# Genomic reconstruction of the successful establishment of a feralized bovine population on the subantarctic island of Amsterdam

**DOI:** 10.1101/2023.11.24.568563

**Authors:** Mathieu Gautier, Thierry Micol, Louise Camus, Katayoun Moazami-Goudarzi, Michel Naves, Elise Guéret, Stefan Engelen, François Colas, Laurence Flori, Tom Druet

## Abstract

The feral cattle of the subantarctic island of Amsterdam provide an outstanding case study of a large mammalian population that was established by a handful of founders and thrived within a few generations in a seemingly inhospitable environment. Here, we investigated the genetic history and composition of this population using genotyping and sequencing data. Our inference showed an intense but brief founding bottleneck around the late 19^th^ century and revealed contributions from European taurine and Indian Ocean zebu in the founder ancestry. Comparative analysis of whole genome sequences further revealed a moderate reduction in genetic diversity despite high levels of inbreeding. The brief and intense bottleneck was associated with high levels of drift, a flattening of the site frequency spectrum and a slight relaxation of purifying selection on mildly deleterious variants. However, we did not observe any significant purge of highly deleterious variants.

Interestingly, the population’s success in the harsh environment can be attributed to pre-adaptation from their European taurine ancestry, suggesting no strong bioclimatic challenge, and also contradicting evidence for insular dwarfism. Genome scan for footprints of selection uncovered a majority of candidate genes related to nervous system function, likely reflecting rapid feralization driven by behavioral changes and complex social restructuring. This unprecedented case study provides valuable insights into rapid population establishment, feralization, and genetic adaptation in challenging environments. It highlights the importance of preserving the unique genetic legacies of feral populations and raises ethical questions in the eyes of conservation efforts.

## Introduction

Understanding the successful establishment of introduced animal species in novel environments is a fundamental area of inquiry in ecological and evolutionary research. The criteria for a population to be deemed “established” involve its capacity for self-sustained reproduction without additional introductions (Sol, 2007). The success of this establishment is contingent upon various factors, including the adaptability of non-native species to the new environment. Among these factors, the overall number of founders and their spatio-temporal patterns of arrival that define the “propagule pressure” plays a pivotal role (Simberloff, 2009). Both empirical and theoretical evi-dence demonstrates a positive relationship between propagule pressure and establishment probability (Lockwood *et al*., 2005; Simberloff, 2009; Cassey *et al*., 2018). Indeed, higher propagule pressure typically involves more in-dividuals being introduced, potentially carrying diverse genetic backgrounds thereby increasing genetic variation within the founding population which can then adapt to a wider range of challenges and enhance the population’s chances of survival. Conversely, this makes it challenging for a small number of individuals (<10) to establish in a non-native habitat (Cassey *et al*., 2018).

However, this conventional understanding of genetic diversity’s importance is challenged by the “genetic paradox of invasion.” This paradox arises from numerous cases of non-native species becoming invasive with only a small founding population (Estoup *et al*., 2016). It can however be resolved if there is no substantial genetic impover-ishment in the invasive populations compared to the native ones, or if the founding individuals were pre-adapted to the new environment, as in the scenario known as “anthropogenetically-induced adaptation to invade” (AIAI) (Hufbauer *et al*., 2012; Estoup *et al*., 2016). Additionally, bottlenecks can facilitate population survival by inten-sifying the exposure of deleterious mutations in homozygous individuals. Such purging of genetic load is usually considered to be more effective for bottleneck of intermediate intensity (e.g., moderate reductions in population size over several generations) and for highly deleterious and recessive variants (Glémin, 2003; Estoup *et al*., 2016). Finally, fast life histories, which are characterized by frequent reproduction, high fecundity, and early maturity, that may lead to rapid population growth are typically viewed as reducing the risk of extinction (Capellini *et al*., 2015). However, in several species including mammals, the link between fast life histories and establishment success was found to be inconsistent (Sol *et al*., 2008; Capellini *et al*., 2015).

The cattle population on the sub-antarctic island of Amsterdam, a secluded 55 km^2^ island located 4,440 km south-east of Madagascar, presents a remarkable case of successful establishment. This population quickly thrived in an apparently inhospitable environment and despite founded by only a handful of introduced bovines. Indeed, historical records suggest that it was founded by five (or six) individuals brought to the island in 1871 by a farmer from La Réunion island named Heurtin (Lesel *et al*., 1969). He and his family attempted to settle on the island but ultimately found the living conditions unbearable, leading to their departure a few months after arrival and the abandonment of their cattle. Some alternative hypotheses propose that Heurtin relied on cattle already introduced by navigators throughout the 18^th^ and early 19^th^ centuries to secure a source of meat for future journeys since at that time, Amsterdam Island served as an important landmark for transoceanic travelers and sealers (Lesel *et al*., 1969). Nonetheless, there are no documented instances of cattle sightings before 1871 (in contrast to other livestock such as pigs or goats which have never established) to support these claims (Micol and Jouventin, 1995). From 1871 onwards, the island’s cattle population lived freely and multiplied, reaching its peak of around 2,000 individuals in 1952 and in 1988 (Micol and Jouventin, 1995), and becoming one of the few known feral cattle populations (Hall, 2008).

Despite being threatened by a suspected paratuberculosis outbreak in 1953 that led to a sharp decline to 800 heads (Fiasson *et al*., 1953; Lesel *et al*., 1969), the population quickly recovered. However, as the proliferation of cattle was seen as a threat to the endemic woody shrub related to the *Phylica arborea* species and to the endemic alba-tross, *Diomedea amsterdamensis*, control measures were implemented in 1988. The size of the cattle population was reduced to around 500 in 1993, and two long fences were erected, enclosing the animals within a 1225-acre area (Micol and Jouventin, 1995). The authorities opted finally for a full eradication of the bovine population in 2010, ignoring protests from several scientists and the possibility of coexisting with appropriate population man-agement policies and control measures against the endemic albatross’ direct predators, including cats, rats, and mice. Additionally, no endeavor was made to gather samples, such as blood, tissue, or semen, for subsequent sci-entific analysis. Nevertheless, we were able to recover DNA samples of 18 individuals obtained from two previous sampling campaigns in 1992 and 2006. These samples were sufficiently preserved to be genotyped, and for eight of them, whole genome sequencing (WGS) was conducted.

This study aims to provide a comprehensive genomic characterization of Amsterdam island cattle population, re-ferred to as TAF population hereafter, based on these newly generated data. The goal is to gain insights into the genetic foundation that enabled the successful establishment and feralization of a large domestic mammal from an extreme but short population bottleneck. Our analysis first includes a detailed inference of demographic history based on genetic data. Second, we relied on WGS data to assess levels of genetic diversity and genetic load, thereby looking for evidence of relaxation of purifying selection or purging of deleterious variants. Finally, we jointly used WGS and genotyping data to characterize the adaptive history of the TAF population, including i) an assessment of the genetic maladaptation (Capblancq *et al*., 2020) of domestic cattle to the Amsterdam island environment; ii) the critical evaluation of the island dwarfism hypothesis as recently proposed based on crude phenotypic data (Rozzi and Lomolino, 2017); and iii) the identification of key physiological functions mobilized by selection from the genes surrounding our identified genomic footprints of selection.

## Results

### The genetic history of the cattle of the Amsterdam island

A total of 18 TAF individuals (Figure 1A) sampled in 1992 (n=12) and 2006 (n=6) during two different campaigns (Table 1) were newly genotyped on the commercial BovineSNP50K assay (Matukumalli *et al*., 2009) along with 31 individuals from the Moka Zebu (MOK) population, a local breed living on the La Réunion island that was considered a possible proxy for the TAF founders. These data were combined with public data from 30 other cattle populations representing the worldwide cattle diversity including, i) two Brazilian Zebu breeds originating from India (ZEB); ii) 3 African taurine (AFT) breeds; iii) 4 African Zebu breeds (AFZ) including the two closely related Indian Ocean Zebu breeds from the islands of Mayotte (MAY) and Madagascar (Magnier *et al*., 2022); and iv) 21 European (mostly French) taurine breeds (EUT) (Table 1, Figure 1B). After filtering, the final combined dataset (hereafter referred to as W50K), consisted of 876 individuals from 32 different populations genotyped at 40,426 autosomal SNPs.

**Figure 1.**
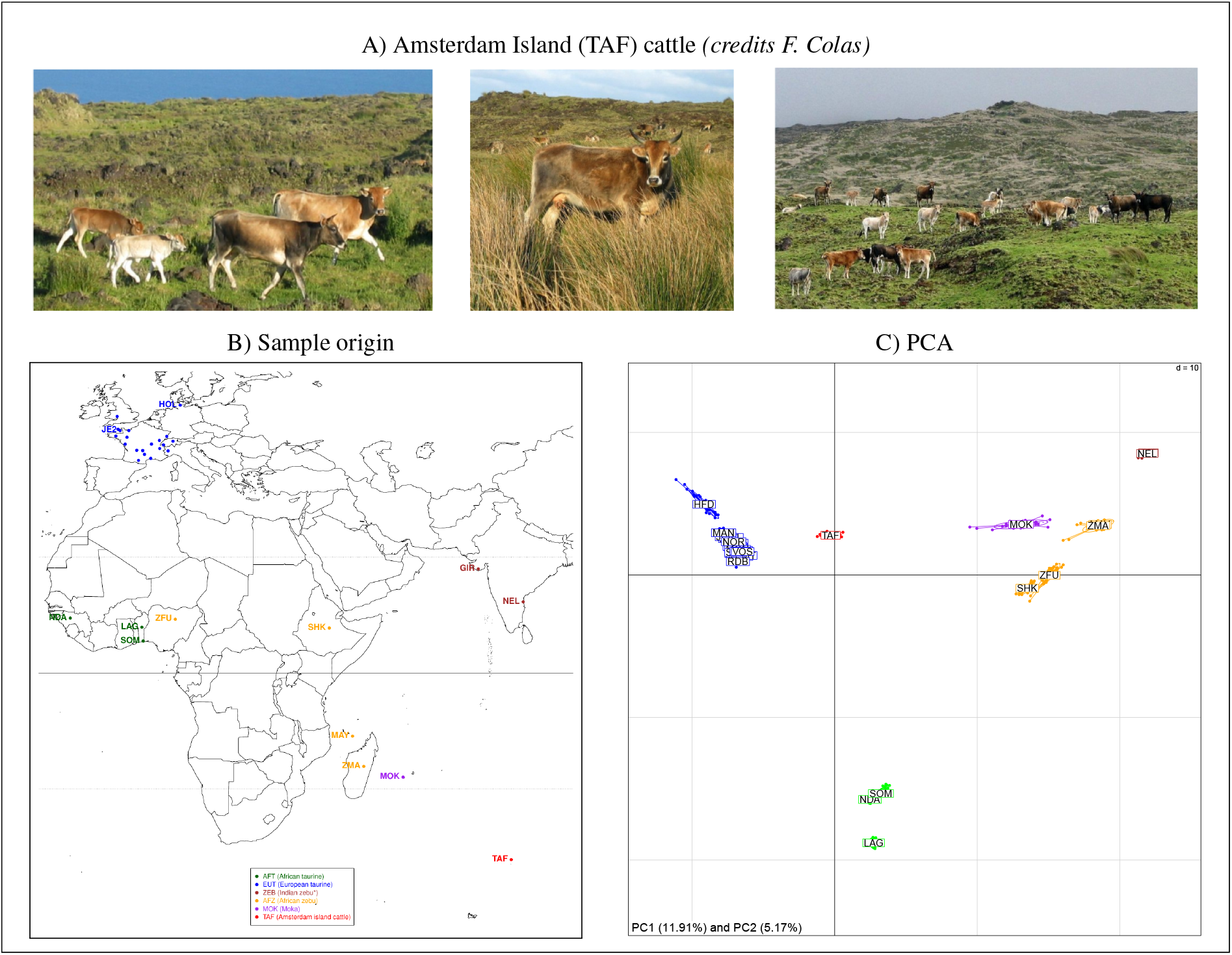
A. Pictures of cattle from the island of Amsterdam island (credits:François Colas), B. Location map of cattle breeds included in the W50K dataset, C. Results of the principal component analysis of the W50K dataset. Individuals are plotted on the first two principal components according to their coordinates. Ellipses characterize the dispersion of each breed around its centroid.

**Table 1.**
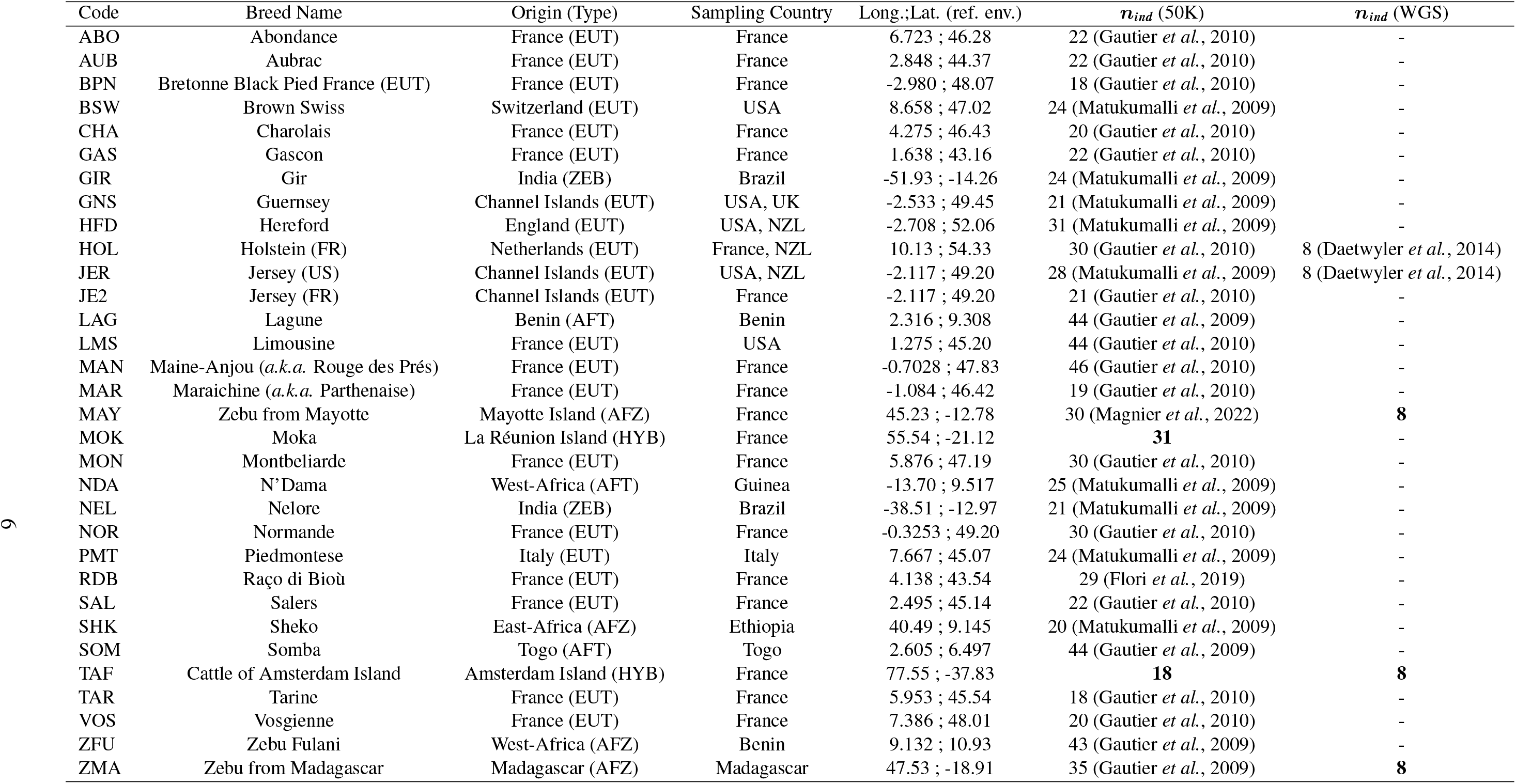
Sample description with country of origin and population type (i.e. EUT=European Taurines, AFT=African Taurines, AFZ=African Zebus, ZEB=Zebus of Indian origin, or HYB=Hybrids); the country of sampling; the GPS coordinate (in decimal degrees) used to retrieve environmental covariates; the number of individuals genotyped on the BovineSNP50 Illumina assay and included in the W50K dataset; and the number of individuals sequenced (WGS) with associated reference. GPS coordinates specifying the environment of each population were taken as the birthplace of the breed including for GIR and NEL in Brazil (Gautier *et al*., 2010). Individuals from the Jersey (EUT) breed were divided into two distinct groups corresponding to i) 21 individuals (coded JE2) sampled in northwestern France (Maine-et-Loir county) and born between 1982 and 1990 (Gautier *et al*., 2010); and ii) 28 individuals (coded JER) sampled in the USA (Matukumalli *et al*., 2009). The newly genotyped (n=49) and sequenced (n=24) individuals are highlighted in bold. All genotyped data (including the new ones) are available from the WIDDE repository (Sempéré *et al*., 2015) and the newly generated whole genome sequence data are publicly available in the NCBI SRA repository (project PRJNA1010533).

#### Genetic structuring of the TAF population

As detailed in Table S2, 43.3% of these SNPs were monomorphic in the TAF sample which was the highest level among observed values ranging from 39.6% to 40.8% in ZEB; 23.5% to 41.0% in AFT; 19.5% to 34.0% in AFZ; and 7.41% to 21.5% in EUT. Due to the strong SNP ascertainment bias toward SNPs of EUT origin (Gautier *et al*., 2010; Matukumalli *et al*., 2009), the within-population genetic diversity estimates obtained from the BovineSNP50K assay data must be interpreted with great caution, particularly for the ZEB, AFZ and AFT breeds (see below for accurate estimates based on WGS data). Nevertheless, the estimated heterozygosity of 0.216 observed in TAF was lower than that observed in EUT (from 0.258 in RDB to 0.323 in HFD) and in the range of that observed in AFZ populations (from 0.204 in ZMA to 0.255 in SHK). More interestingly, the *F_IS_* estimated within the TAF sample was found to be significantly negative indicating an excess of heterozygotes in the sample relative to the Hardy-Weinberg equilibrium expectation, although the 18 samples from the 1992 and 2006 campaigns were analyzed together. In addition, the estimated pairwise kinship coefficients among the 12 samples collected in 1992 (Manichaikul *et al*., 2010) ranged from-0.039 to 0.26 (median of 0.014), suggesting some close parental relationships (Figure S1). For example, the relationship between TAF 5609 (born 1985) and TAF 5608 (born 1989) was estimated to be within [0.177, 0.354], suggesting a first-degree relationship corresponding to a mother-daughter relationship according to the sharing of IBD (Identical by descent) segments. The four coefficients between TAF 5608 and TAF 5612 (*ϕ* = 0.144); TAF 5610 and TAF 5617 (*ϕ* = 0.131); TAF 5609 and TAF 5619 (*ϕ* = 0.101) and TAF 5608 and TAF 5610 (*ϕ* = 0.0887) were within [0.0884, 0.177], suggesting a second-degree relationship. Analysis of IBD segment sharing allowed the relationship between TAF 5608 and TAF 5612 to be upgraded to a full-sib relationship, birth dates suggesting half sibs or aunt-niece relationships for the other three pairs. Note that at least four of these six females belonged to a same group of about 40 individuals. The remaining 61 coefficients between the 1992 individuals were all < 0.0884 (relationship more distant than second degree). Similarly, among the 15 pairs of 2006 samples, the estimated kinship coefficients were < 0.0884 ranging from-0.035 to 0.074 (median of 0.013), as were the 72 pairwise comparisons between individuals collected in 1992 and 2006 which ranged from-0.091 to 0.062 (median of-0.0081) and were shifted toward lower values, as expected.

#### Relationship with other worldwide populations

The global *F_S T_* estimated on the W50K dataset was found to be 21.5% (CI95%=[21.1%; 21.8%]) while the global *F_IS_* remained negligible (CI95%=[0.04%; 0.26%]). As shown in Figure S2, the estimated pairwise *F_S T_* values ranged from 0.0247 for the MAY/ZMA pair to 0.472 for the LAG/NEL pair (median of 0.203). For pairs including the TAF, values ranged from 0.208 (TAF/PMT) to 0.436 (TAF/NEL), consistent with a presumably high amount of drift in this population following an intense founding bottleneck. In addition, the TAF appeared more closely related to breeds of EUT origin. However, clustering based on pairwise *F_S T_* must be interpreted cautiously and cannot be safely used to infer population origins as it is highly sensitive to drift and admixture. For example, the TAF may appear as an outgroup of the EUT cluster in Figure S2 due to drift and the TAF/PMT pair is likely to display the lowest pairwise *F_S T_* because, like the TAF, the PMT breed has some amount of ZEB ancestry (see Figure S4 and Gautier *et al*., 2010). Finally, note that the *F_S T_* between the TAF and the newly genotyped MOK sample was 0.275, i.e., in the upper range of values estimated for pairs involving this population sample, which varied from 0.0578 for the MOK/ZMA pair to 0.303 for the MOK/LAG pair.

To refine the exploration of the overall structuring of genetic diversity on the W50K dataset, we computed allele-sharing distances between all pairs of individuals and used the resulting matrix to build the neighbor-joining tree shown in Figure S3. In agreement with previous observations (e.g. Gautier *et al*., 2010) and the results above, the obtained tree showed a clear clustering of individuals according to their population of origin, except to some extent for the MOK individuals, which were more dispersed with two apparent outliers. At a higher level, the EUT, AFT and ZEB populations were also well separated, as were the AFZ and MOK populations, both of which branched between ZEB and AFT. Finally, the TAF population was clearly isolated from the other populations and branched close to the EUT populations. A similar pattern was recovered using principal component analysis (PCA) of the W50K dataset (Figure 1C). Interestingly, in the first factorial plan, TAF and MOK individuals are positioned on an axis connecting the ZMA and EUT populations, suggesting ZEB and EUT ancestry for both populations, with TAF individuals closer to EUT and MOK closer to ZMA and MAY. An alternative but qualitatively similar representa-tion was obtained using unsupervised hierarchical clustering (Figure S4). For example, at K=3 predefined clusters, the proportion of the “green” cluster (interpreted as EUT) ranged from 68.0% to 75.0% (median of 71.4%) for TAF individuals and from 19.8% to 26.4% (median of 23.0%) for MOK individuals. However, the exact origins of the different ancestries remain difficult to assess with such exploratory analyses and should therefore be interpreted with caution.

#### Inferring the demographic history of the TAF population

We relied on the *f*-statistics based framework (Patterson *et al*., 2012) as implemented in the R package poolfstat (Gautier *et al*., 2022) to infer the origin of the TAF population using the W50K dataset. We first performed a formal test of admixture based on the *F*_3_ statistic computed for all the 465 population triplets with TAF as the target admixed population, but none of the resulting statistics were found to be (significantly) negative. In fact, this can be explained by the high amount of drift due to the extreme founding effect in the TAF history. Conversely, 100 of the 465 population triplets with MOK as the target population showed significantly negative *f*_3_ (at the 5% threshold), with the lowest value observed for the (MOK;GIR,NDA) and (MOK;BPN,ZMA) configurations (both with *Z* = −21.7).

We also used *f*-statistics to construct an admixture graph construction as detailed in Figure S5 (Patterson *et al*., 2012; Gautier *et al*., 2022). This allowed us to demonstrate the admixed origin of the TAF and MOK populations with closely related EUT ancestral sources contributing to 75% and 22% of their genome composition, respectively, and another source closely related to ZMA. Positioning the MOK population resulted in a poorer fit suggesting a more complex origin (i.e., three-way EUTxZMAxZEB admixture) difficult to capture with the W50K dataset (Figure S5). Nevertheless, the inferred graph suggests that MOK may not be considered as the closest proxy for the source populations from which TAF originated. Figure 2A shows the results for the best-fitting graph among all possible ways of positioning TAF and the JE2 Jersiaise breed sample (Table 1) on a scaffold graph previously inferred to describe the history of the Indian Ocean island populations MAY and ZMA (Magnier *et al*., 2022). This graph included, in addition to MAY and ZMA, two ZEB (GIR and NEL), two AFT (LAG and NDA), an East African Zebu (here replaced by the closely related SHK) and an EUT breed (HOL). The graph was actually very similar to that of Figure S5B. Note that we chose to consider JE2 because it was the EUT breed the most divergent to HOL (Figure S4 with K=4) and provided a very similar admixture-graph than the one obtained using the JER sample but with a slightly better fit (not shown). With the addition of MAY, the ancestral TAF source related to Indian Ocean Zebu branches before the separation of MAY and ZMA, which we previously estimated to have started around the 16^th^ century (Magnier *et al*., 2022). This result suggests that the population contributing to the Indian Ocean Zebu source was established early on the island of La Réunion. Alternatively, more recent exchanges between MAY and ZMA (Magnier *et al*., 2022) may have contributed to their closer proximity on the inferred graph. To provide insight into the EUT origin of the TAF (see Figure S6), we ranked the 22 sampled EUT breeds by their proximity to the EUT ancestral source of the TAF using i) the *f*_3_ estimates for all the (TAF;ZMA,EUT) population triplet configuration (ranked in ascending order, Table S4); and ii) the *f*_4_ estimates for all the (EUT,NDA;TAF,GIR) population quadruplet configuration (ranked in decreasing order, Table S5). Although all confidence intervals (CI) overlap, the closest breeds originate from northwestern Europe, in particular the island of Jersey. Based on the inferred admixture graph, we could obtain a more accurate estimate of the 95% CI for the admixture proportions using *F*_4_-ratio (Figure S6) giving 73.1% (*CI*_95%_ = [70.7; 77.3]) of EUT ancestry and 26.9% (*CI*_95%_ = [22.7; 29.3]) of Indian Ocean Zebu ancestry in the TAF population (consistent with Figure 2A). Interestingly, the eight sequenced TAF individuals (see below) all carried the mitochondrial haplogroup “T1b1” found in most African cattle, including African zebus (Ward *et al*., 2022). This same haplogroup was also present in 6 of the 8 ZMA and one of the 8 MAY individuals sequenced, with the remaining two ZMA and 5 MAY individuals carrying the closely related “T1b1b1a1” haplogroup (two MAY individuals carried “T1d”). This suggests that the TAF founders were mixed with local breeds before being moved to the island of Amsterdam.

**Figure 2.**
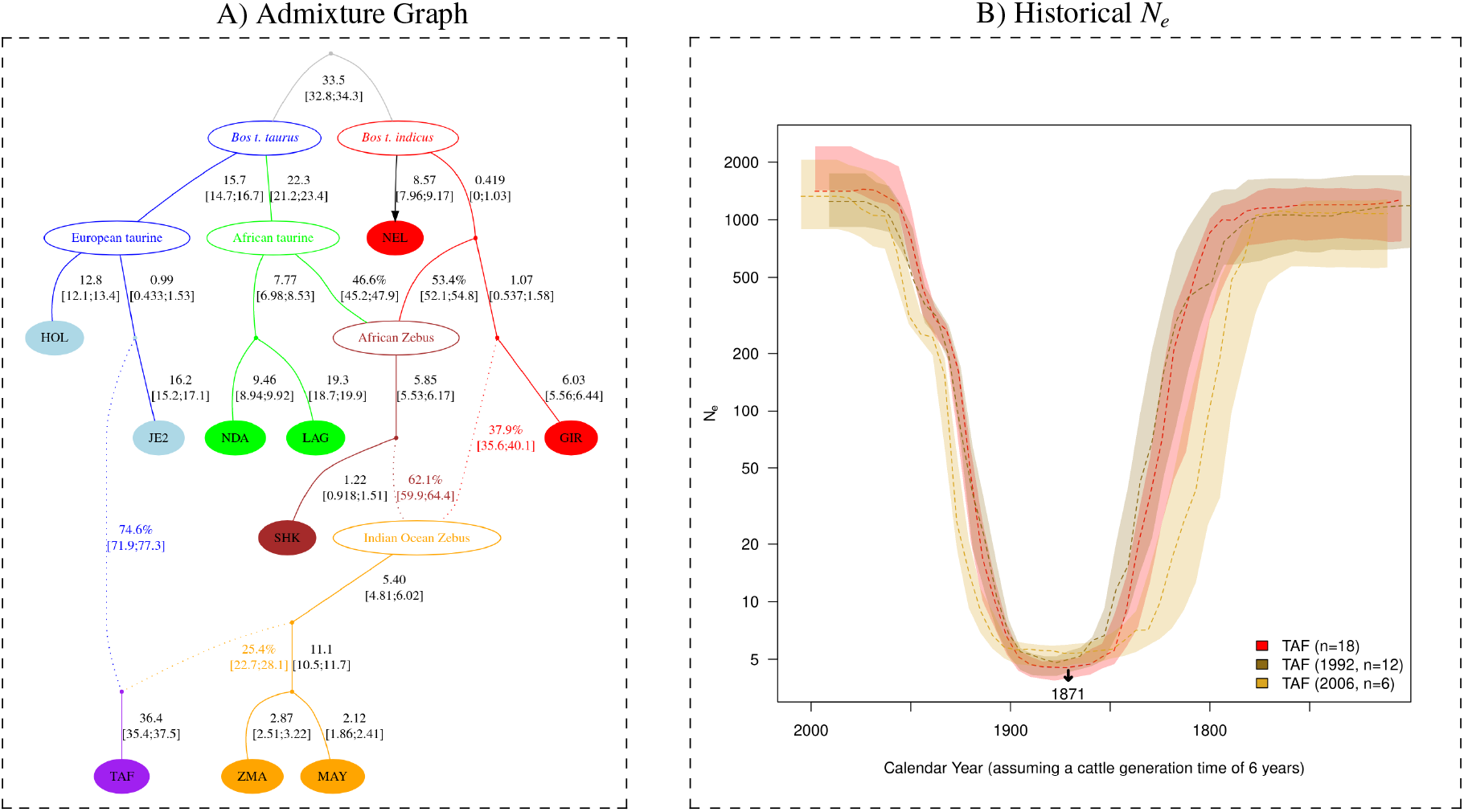
The demographic history of the Amsterdam island cattle (TAF) population inferred from genotyping data (W50K dataset). A) Admixture graph linking the TAF population (in purple) with Indian Ocean Zebu (MAY and ZMA in orange), zebu breeds of Indian origin (GIR and NEL in red), African taurine (NDA and LAG in green) and European taurine breeds (HOL and JE2 in blue) inferred with poolfstat (Gautier et al., 2022). Admixture events are indicated by dotted arrows. Estimates of branch lengths in drift units (x100) and admixture rates are given next to the corresponding edges with (approximate) block-jackknife 95% CI in brackets. This is the graph with the best fit (*BIC* 3.4 units lower than the graph with the second lowest BIC) among all possible ways of positioning TAF and JE2 on a scaffold graph reproducing the graph obtained by (Magnier et al., 2022) to infer the history of Indian Ocean island populations. The Z-score for the worst fitting f-statistics, f4(HOL, JE2; MAY, NDA) is −2.02. B) Recent population size history (Ne) estimated with the program Gone program (Santiago et al., 2020) for the TAF population analyzing the two samples collected in 1992 and 2006 separately and together. For each samples (with size, collection date and color code indicated in the legend box), the average Ne trajectories (dashed line) and 95% confidence envelope estimated from blockjackknife sampling are plotted. The time scale has been transformed to calendar years assuming a 6-year generation time for cattle and accounting for each collection date. The arrow on the x-axis indicates the founding date of the TAF population (1871) according to historical records.

#### Timing of admixture of the TAF source population

We further estimated the timing of this admixture event using the alder program, which is based on modeling the exponential decay of admixture-induced Linkage Disequilibrium (LD) with genetic distance (Loh *et al*., 2013). Using TAF as the target admixed population in one-reference tests (i.e., using an LD measure weighted by allele frequencies observed in a single source population proxy), a significant weighted LD curve was found with all the 31 other breeds of the W50K dataset, except MOK and ZFU, confirming the admixture signal. We further fitted the TAF two-reference weighted LD curves (i.e., using an LD measure weighted by the allele frequencies observed in two source proxies) for all 406 possible pairs of the 29 populations that passed the one-reference tests. A total of 73 tests were considered successful (i.e., gave parameter estimates consistent with the one-reference fitted curve obtained with either of the two reference populations). The highest amplitude (i.e., y-intercept) estimate was obtained with NEL and JE2 as the source proxies suggesting that these populations were the best closest proxy to the actual source populations (among those sampled that passed the tests). The corresponding estimated time for admixture was found to be *t_a_* = 22.02 ± 2.50 generations (i.e., year 1860 ± 15 assuming a 6-year generation time and 1992 as the average birth year) and was consistent across all tests. Note that all two-reference weighted LD curves with ZMA (or MAY) and an EUT breed gave similar results, but the best-fitting estimates of the timing of admixture were discordant between those obtained with fitting the decay of LD weighted with the two or either one of the two references. More specifically, the timing of admixture for the one-reference-weighted LD curve was always significantly lower with ZMA or MAY as the source proxy (*t_a_* = 17.48 ± 2.48 and *t_a_* = 18.26 ± 2.36, respectively) than any other EUT population or ZEB population. Although difficult to prove formally, this may indicate that the TAF founders had heterogeneous EUT ancestry, i.e., one or some of the few founders may have had substantially more Indian Ocean Zebu ancestry than the others.

#### The TAF recent size population history

Figure 2B plots the recent evolution of the effective population size of the TAF population as inferred using Gone (Santiago *et al*., 2020) with the W50K dataset after separating the 1992 (n=12) and 2006 (n=6) samples. The two resulting curves were consistent and both showed a clear bottleneck down to *N_e_* = 4.80 (20 generations ago) and *N_e_* = 5.38 (24 generations ago), respectively, and *N_e_* = 4.52 (22 generations ago) when both samples are combined. Such estimates are remarkably consistent with the historical data referring to 5 founders in 1871. The timing also coincides with the estimated admixture time (22 generations ago), suggesting that admixture occurred almost simultaneously with the founding of the TAF population, consistent with the above hypothesis from the alder results about the possible genetic heterogeneity of the founders. Finally, Figure 2B shows that the bottleneck was followed by a very steep increase in TAF population size toward the final effective size of about 1,500 individuals, although no evidence of a decline corresponding to the paratuberculosis outbreak reported in the 1950s could be found. It should also be noted, that the above analysis of mitochondrial sequences based on the WGS data we analyzed below is consistent with a very small number of introduced founders since only one haplogroup (and only two polymorphic positions) was found among the 8 individuals.

### Characterization of the genetic diversity with WGS data

We were fortunately able to recover enough high-quality DNA of 8 TAF females sampled in 1992 to perform their whole genome sequencing together with 8 ZMA and 8 MAY individuals as representative of their Indian Ocean Zebu ancestry (Table S3). We analyzed these 24 newly generated WGS data with publicly available ones for 8 JER and 8 HOL representing the EUT breeds, choosing data with similar coverage characteristics.

#### Within population genetic diversity

Based on these WGS data, we first characterized the impact of the demographic history on inbreeding and diversity levels in the TAF and compared these with those observed in other populations. Importantly, all these populations have experienced different demographic histories since the establishment of the TAF population. In particular, European cattle have been intensively selected and have undergone a continuous decline of their effective population sizes over recent generations (e.g. Gautier *et al*., 2010), whereas Indian Ocean Zebu populations have experienced more ancient and less intense bottlenecks (Magnier *et al*., 2022).

Despite a very strong founder event, TAF individuals exhibited slightly higher genome-wide heterozygosity than the European breeds JER and HOL, but, as expected, significantly lower than the Indian Ocean Zebus ZMA and MAY (Figure 3A). Similar conclusions could be drawn when comparing the number of segregating sites, which was almost halved in TAF compared to MAY and ZMA, but (slightly) higher than in HOL or JER (Figure 3C and Table S6). Similarly, the ranking in terms of fixation levels was reversed due to increased drift in small populations, with four times more fixed variants in TAF compared to MAY or ZMA Zebu, but more than 20% less compared to HOL and JER (Table S7). A striking difference in the TAF was the unfolded Site Frequency Spectrum (SFS), which was flattened compared to other populations (Figure 3B), with fewer rare alleles and more alleles segregating at higher frequencies. To further characterize the effect of the recent TAF history on the individual levels of inbreeding, we partitioned individual genomes into different classes of autozygous (aka homozygous-by-descent or HBD) segments (Druet and Gautier, 2017). According to our model, the length of HBD segments is assumed to be exponentially distributed with a class-specific rate (i.e., the expected length is specific to each HBD class). Each HBD class thus corresponds to a distinct group of past ancestors (e.g. long segments correspond to a group of recent ancestors). We observed high levels of inbreeding in the TAF population, averaging 33% (Figure 3D). Interestingly, autozygosity was concentrated in the HBD class associated with ancestors present about 15 generations in the past (i.e., the HBD class with rate *R_c_* equal to 32 and an expected length of about 3 cM). This is fully consistent with the founding event and the demographic history estimated under the Gone model (Santiago *et al*., 2020), especially if we consider that neighboring classes corresponding to 8 (*R_c_*= 16) and 32 (*R_c_*= 64) generations respectively are more distant from the founder event. These values were higher than recent levels of inbreeding (*R_c_* ≤ 128) observed in Indian Ocean Zebu (MAY and ZMA) or EUT (HOL and JER) populations, despite intensive selection and reduced effective population size. Strikingly, however, the TAF individuals had few long HBD segments associated with recent ancestors (recent HBD classes had low contributions). This indicates that the population expanded rapidly (the period of reduced *N_e_* was short), consistent with demographic estimates, and that most of the inbreeding occurred a few generations after the bottleneck. In contrast, individuals from the two EUT dairy breeds HOL and JER showed higher levels of recent inbreeding, despite management efforts to avoid it. Finally, we compared the partitioning estimated with the 50K SNP genotyping data between the 12 individuals sampled in 1992 and the 6 individuals sampled in 2006, where the SNP density was sufficient to obtain a good resolution for recent times, up to about class *R_c_* = 128 (representative of ancestors living 64 generations ago), i.e., before the establishment of the TAF population (Druet and Gautier, 2017). As shown in Figure S7, autozygosity levels were stable (from 32.6 to 32.9% on average), but concentrated in more ancient HBD classes for the most recent group of individuals (inbreeding levels in the most recent classes *R_c_* ≤ 16, dropped from 5.9 to 2.1% on average), indicating that ancestors contributing to inbreeding are becoming more distant with time. This confirms that new inbreeding is not generated and that autozygosity is mainly related to the founder event.

**Figure 3.**
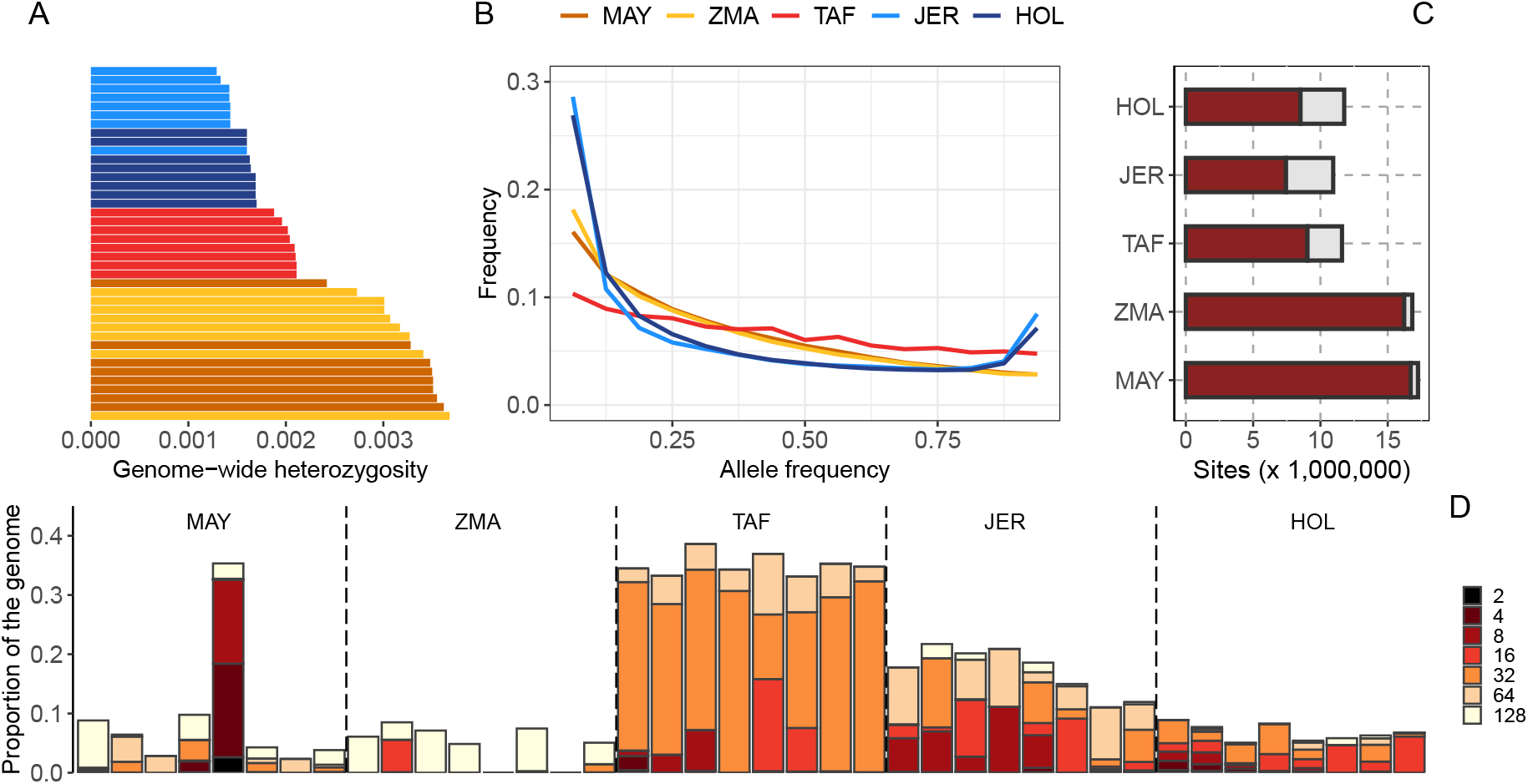
Genetic diversity estimated with WGS data in the TAF populations and populations representative of its European taurine (JER and HOL) and Indian Ocean Zebus (ZMA and MAY) ancestry. A) Genome-wide nucleotide diversity (heterozygosity) of the 40 individuals ranked in increasing order and colored according to their population of origin. B) Site-frequency spectrum estimated for the five breeds. C) Number of sites per population that are polymorphic (brown) and fixed (grey) for the derived allele. D) Inbreeding levels, and partitioning of inbreeding in different HBD classes in the different populations.

#### Genetic load

Next, we examined the effect of the founder event on the distribution of deleterious variants to investigate whether purifying selection was relaxed or whether some purging occurred as in other species that experienced a severe bottleneck, such as mountain gorillas (Xue *et al*., 2015) or Alpine ibex (Grossen *et al*., 2020). To this end, we first compared the proportions of variants in different functional categories as in Grossen *et al*. (2020). The TAF was found to have about 5% higher proportions of polymorphic NS (Non-Synonymous) variants than MAY and ZMA, but about 10% lower than European cattle (Table S6). The higher proportion of NS variants compared to the two Indian Ocean Zebus suggests that there has been some relaxation of purifying selection, although the proportions of the most deleterious class of variants (i.e., deleterious NS and Loss-of-Function or LoF) among the segregating sites were always lower (from 1% to 28%) in TAF than in the other four populations. To investigate this further, we compared the Site Frequency Spectrum (SFS) for the different classes of variants (Figure 4A-C). In TAF, the curves were flattened for all categories, whereas in the other populations, the SFS calculated for deleterious variants were enriched in the low frequency classes (Figure S8). Consequently, there are proportionally fewer NS or LoF variants with frequency > 0.25 in the control populations (JER, HOL, MAY and ZMA), whereas in TAF the proportion of NS and LoF functions is close to that observed for other variant classes (including intergenic variants) up to allele frequencies close to 75% where it starts to decrease. This suggests that NS and LoF variants can reach higher frequencies more often in TAF.

**Figure 4.**
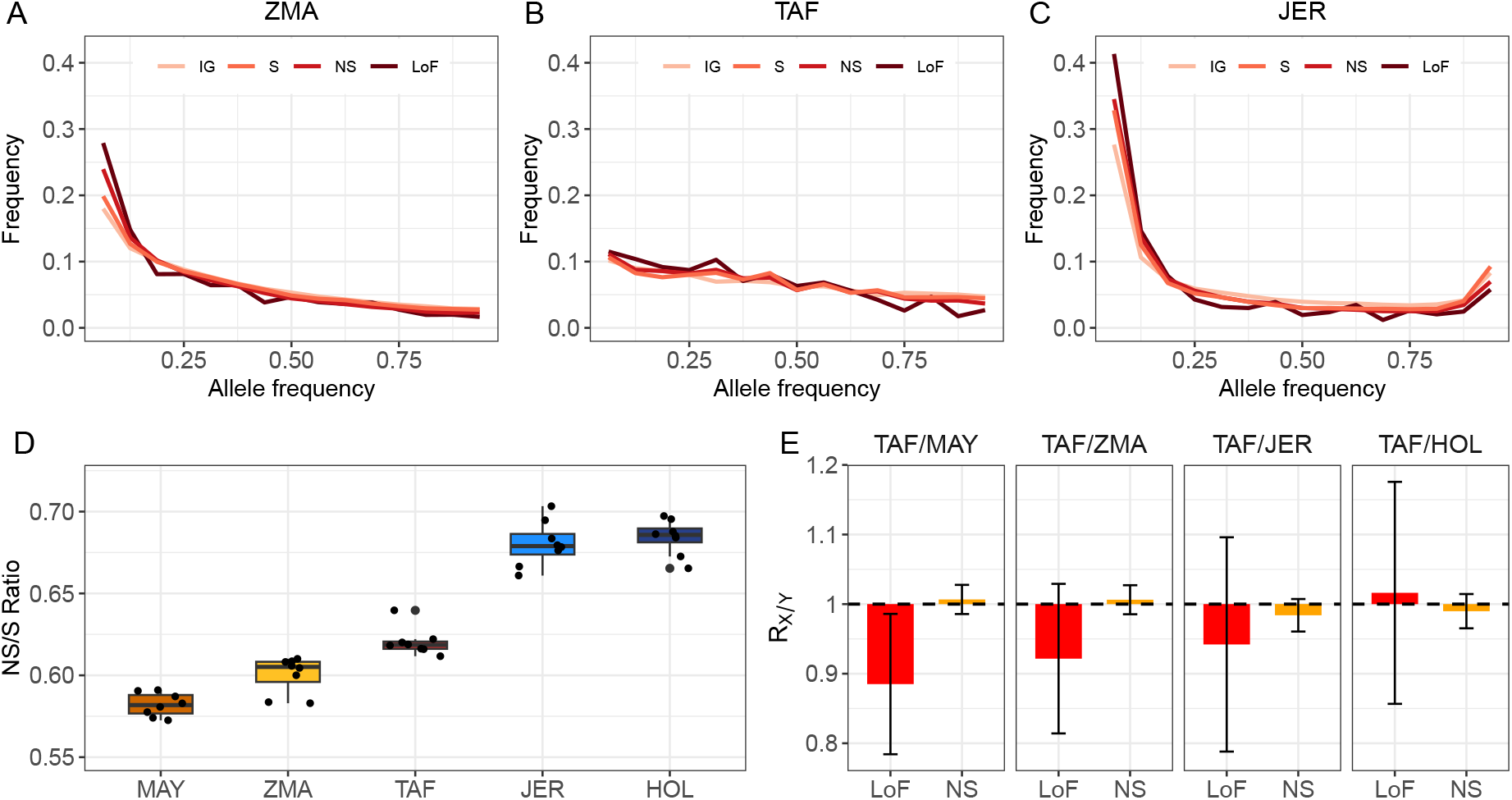
Genetic load in cattle populations. A-C. Comparison of the site frequency spectrum for intergenic (IG), synonymous (S), non-synonymous (NS) and loss-of-function (LoF) variants in the three breeds. D. Ratio of heterozygous NS versus S genotypes per individual. E. *R_X_*_/*Y*_ statistics for comparisons between TAF and other breeds for NS and LoF variants.

The ratio of the number of heterozygous NS to synonymous (S) genotypes per individual has previously been used as evidence for increased mutation load, for example as a result of domestication (Cruz *et al*., 2008; Renaut and Rieseberg, 2015). We observed that this ratio was only slightly higher in TAF than in MAY and ZMA (Figure 4D) but remained significantly lower than in JER and HOL, the two intensively selected dairy cattle breeds, suggest-ing some relaxation of selective constraints in TAF. Next, following Smeds and Ellegren (2023), we divided the genetic load into masked and realized load, estimated as the number of heterozygous and fixed derived variants per individual, respectively. The TAF population showed a slightly higher masked load than HOL and JER for all classes of variants, but it remained well below the levels observed in MAY and ZMA (Figure S9). The trends were reversed for the realized load, which may be the result of the high inbreeding levels and fixation rate observed in TAF (and HOL and JER) compared to MAY and ZMA. In general, the number of homozygous genotypes (realized load) is more relevant when deleterious effects are recessive, whereas the number of derived alleles per genome would be more relevant for additive effects and is predicted to be insensitive to demographic history for neutral alleles (Mooney *et al*., 2023). Interestingly, we observed that the frequencies of derived intergenic variants in our populations were all very similar, ranging from 0.277 to 0.279 (with three populations at 0.278), consistent with expectations for neutral alleles. This suggests that our data processing, including variant filtering and polarization, was implemented reliably (Mooney *et al*., 2023). We therefore confidently used the allele frequencies to compute the *R_X_*_/*Y*_ statistic (Do *et al*., 2015), a relative measure that indicates whether deleterious variants in population *X* are under relaxed (*R_X_*_/*Y*_ > 1) or stronger (*R_X_*_/*Y*_ < 1) purifying selection with respect to *Y*. As shown in Figure 4E and in agreement with the proportions of NS variants in the different populations (Table S6) and other statistics described above, *R_T AF_*_/*Y*_ was slightly > 1 for comparisons with Indian Ocean Zebus (Y=MAY and Y=ZMA) and < 1 for comparisons with EUT populations (Y=HOL and Y=JER), although the value of 1 could not be excluded from the (block-jackknife 95% CI in all comparisons. For LoF variants (Figure 4E), *R_T AF_*_/*Y*_ were < 1 for all com-parisons except for Y=HOL (but only significant for Y=MAY). This suggests a slightly higher level of purging for this category of variants in TAF.

### Adaptive history of the TAF population

#### Genetic O**ff**set

In order to assess the potential environmental challenges that the bioclimatic condition of Amsterdam Island may represent for different (adapted) cattle populations originating from different regions, i.e. their relative maladap-tation or pre-adaptation, we used the Genetic Offset (GO) statistic (Capblancq *et al*., 2020). For this purpose, we relied on the BayPass GEA model (Gautier, 2015) to summarize the relationship between seven bioclimatic PCs (resulting from a PCA of 19 bioclimatic covariates) and the genomic composition of the 32 populations represented in the W50K dataset. We then estimated the GO between the Amsterdam island environment and the birthplace environment of each of the 31 (non-TAF) populations (Figure 5A), following Gain *et al*. (2023). Interestingly, the lowest GO was obtained for the environment associated with the birthplace of JE2 (and JER), which was also the one with the smallest environmental distance, i.e. calculated without considering the association with the bovine (adaptive) genomic composition (Figure 5). The geographically close environment of the island of Guernsey and Normandy that were associated with the birthplaces GNS and NOR also led to small GO, only 12.8% and 20.1% higher than JE2 and JER respectively. Figure 5B further describes the distribution of GO over European environments with respect to the Amsterdam island. Within this broader range, the island of Jersey was also found to be among the regions with the lowest GO, along with the coastal regions of Brittany, Great Britain, Ireland and northern Spain. As expected GO was highest for environments associated with Zebu or African cattle originating from tropical or subtropical regions (Figure 5A) but also for southern and continental European environments (Figure 5B). Overall, this suggests that the climatic conditions on the island of Jersey (or closely related regions) and those on the island of Amsterdam were not so different, and the populations that originated from this region, and that according to demographic inference contributed the most to the ancestry of the TAF population, may have been largely pre-adapted.

**Figure 5.**
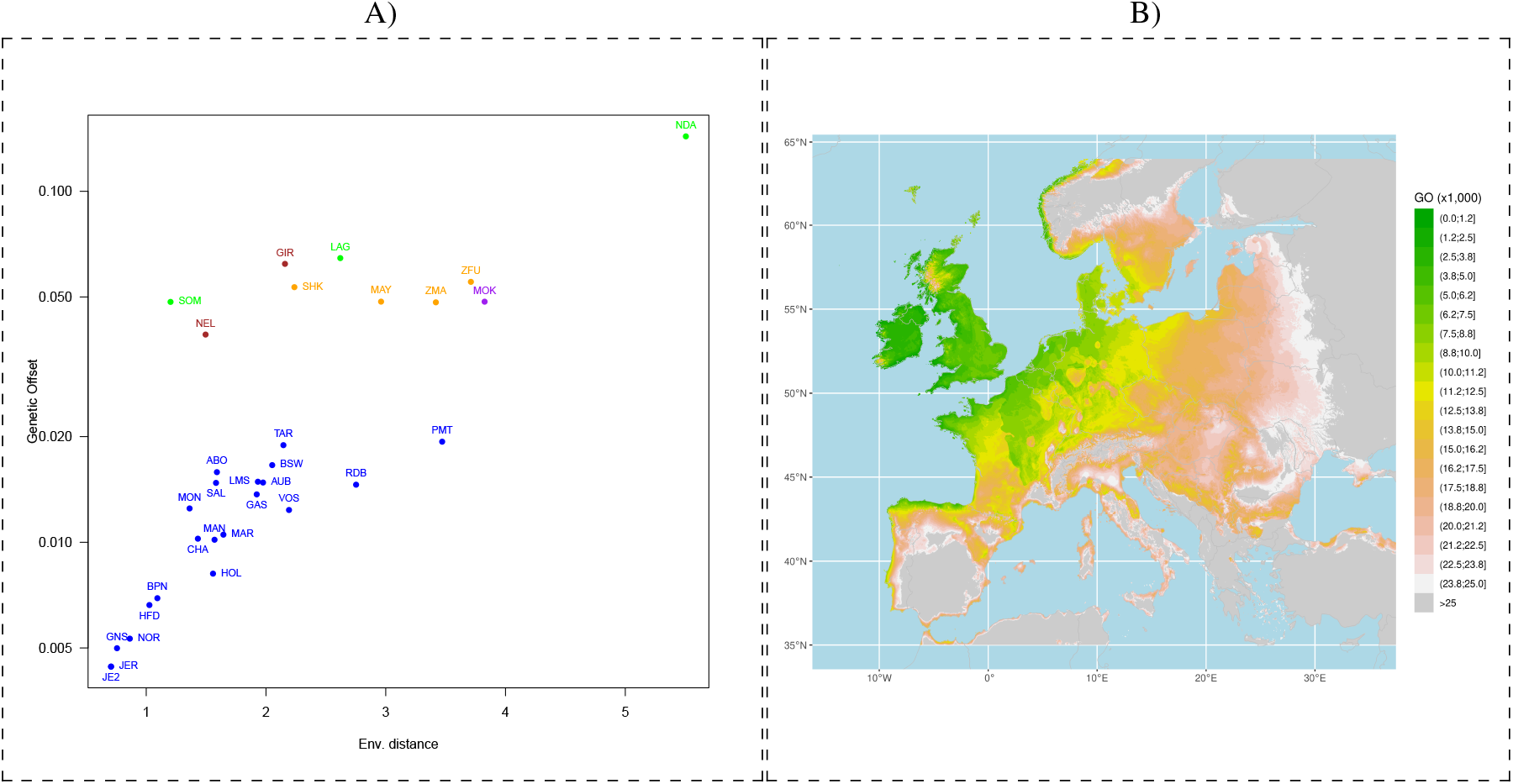
Estimation of domestic bovine maladaptation to the Amsterdam island environment using Genetic Offset (GO) statistics. A) Estimated GO between birthplace environments of 31 populations from the W50K dataset as a function of environmental distance. B) Estimated GO across Europe.

#### Genomic prediction of adult height reveals no evidence for dwarfism in TAF

Although adaptation to the new environment did not appear to be a major challenge, we looked for evidence of selection in the TAF population. For example, based on scarce phenotyping data, Rozzi and Lomolino (2017) argued for rapid dwarfism in this population. If true, this process would have taken place in only a few generations and might have left some strong genomic signatures of selection, at least for the major genes controlling bovine height which would have been polymorphic among the founders. However, our demographic inference showed that the TAF founders are closely related to two populations with short stature (Madagascar Zebu and Jersey individuals). Thus, they may not have been dwarfed as suggested, since the dimensions reported in Rozzi and Lomolino (2017) for TAF cattle are actually consistent with those of the Zebu and Jersey populations, and the TAF would only be short compared to larger cattle breeds (e.g., Holstein). To confirm this, we investigated whether known alleles contributing to short stature were enriched in the population using 164 variants from a recent cattle meta-analysis (Bouwman *et al*., 2018). To do this, we used their estimated genetic effects to predict the height of the sequenced individuals. This amounts to weighting each allele by its predicted effect on stature. We were able to find 105 of these variants in our sequenced individuals (8 per population). As shown in Figure 6, the mean breeding value for (standardized) height was equal to-1.34 for the TAF individuals (ranging from-1.62 to-0.85), and was found to be intermediate between those observed in ZMA (mean of-1.47 with values ranging from-2.16 to-1.28) and in JER (mean of-1.01 with values ranging from-1.73 to-0.34), and substantially lower than for the HOL breed. Accordingly, we observed that the short allele of the PLAG1 mutation (Karim *et al*., 2011) was fixed in all TAF but also in all JER individuals. Thus, there is no evidence that the small size of the TAF individuals is due to (polygenic) selection for “short” alleles.

**Figure 6.**
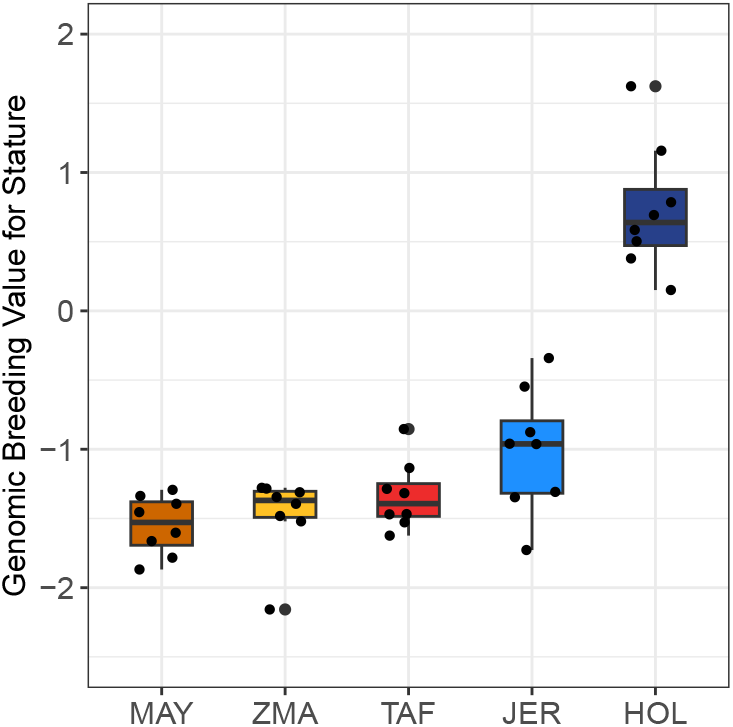
Distribution of genomic prediction of individual height breeding values for each population based on the estimated effects of 105 known variants affecting stature in cattle (Bouwman et al., 2018) genotyped with our analyzed WGS data (n=8 individuals per population).

#### Detecting footprints of selection on a dense haplotype data set

Finally, to provide a global genomic description of the adaptive response of the TAF population in its newly colonized environment, we searched for footprints of positive selection using Extended Haplotype Homozygosity (EHH) based tests, considering either the *iHS* (Voight *et al*., 2006) or the *Rsb* (Tang *et al*., 2007) statistics for within-population and pairwise comparisons, respectively. The *iHS* and *Rsb* based tests are complementary and serve different purposes, the *iHS* being better suitable for detecting recent and strong selective sweeps where the underlying variants are still segregating in the population, while the *Rsb* is more focused on detecting localized and potentially older (but still strong) selective events that lead to fixation of the causal variant in one or the other population. For *Rsb*, we focused here on TAF-specific signals by searching for extended site-specific homozygosity in the TAF with respect to either JER (*Rsb_T AF_*_/*JER*_) or ZMA (*Rsb_T AF_*_/*ZMA*_), considered as source population proxies, using unilateral tests of extremely positive values. To improve mapping resolution, we analyzed a dataset combining high-density (>770,000 SNPs) SNP genotyping data from the Illumina BovineHD assay for 23 ZMA and 30 JER individuals (Magnier *et al*., 2022; Sempéré *et al*., 2015) with the subset of 50K SNP data (most of which are shared with the BovineHD assay) for the 18 TAF individuals. We then imputed (or directly genotyped for the 8 sequenced TAF individuals) the missing BovineHD genotypes in the TAF using the available WGS data (see M&M section). After filtering and phasing, the analyzed data set consisted of 142 haplotypes (36 TAF, 46 ZMA and 60 JER) per autosome, totaling 530,769 SNPs (ranging from 8,842 to 33,710 for chromosomes 25 and 1, respectively). In total we could identify 21 significant regions, 12 (ranging from 76.90 to 1,355 kb in length) associated with the within TAF *iHS*, and two and eight with the *Rsb_T AF_*_/*JER*_ (83.10 and 297.7 kb in length) and *Rsb_T AF_*_/*ZMA*_ (ranging from 196.7 to 1,352 kb), respectively (Table 2). Note that two regions, identified with *iHS* and *Rsb_T AF_*_/*JER*_, overlapped and led to the identification of the same candidate genes (based on the distance to the peak statistic). More signals were identified with *Rsb_T AF_*_/*ZMA*_ than *Rsb_T AF_*_/*JER*_, which may be related to the lower recent historical *N_e_* in JER compared to ZMA, resulting in higher EHH thereby limiting the detection power. In addition, we noticed that two *Rsb_T AF_*_/*ZMA*_ regions overlapped with those detected based on *Rsb_JER_*_/*ZMA*_ statistics (Table S8 and Figure S10). For one of these regions, the identified candidates (and corresponding statistic peaks) were more than 1 Mb apart. We therefore considered them to be separate signals. However, for the other region, the two windows pinpointed the same candidate gene (SLC4A4), suggesting that this region had already been under selection before the establishment of the TAF population (in the ancestral JER-related population) and this non-specific TAF signal was further disregarded.

**Table 2.** Description of the regions containing footprints of selection based on the *iHS*, *Rsb_T AF_*_/*JER*_ and *Rsb_T AF_*_/*ZMA*_ EHH-based tests. The two non-specific TAF regions (#14 and #18) that overlap with those identified using *Rsb_JER_*_/*ZMA*_ (Table S8) are in italics and highlighted with a ^✶^.

| ID | Test | Win. Pos.<br>(chr:start-end in kb) | Win Size in kb<br>(NsnP) | Peak Pos. in kb<br>(Stat. Value) | Nearest Gene<br>(Pos. in kb ; dist. from peak) |
| --- | --- | --- | --- | --- | --- |
| 1 | *iHS*<sub>TAF</sub> | 3:43,306-43,463 | 156.7 (19) | 43,343 (3.87) | AGL (43,352-43,353 ; 8) |
| 2 | *iHS*<sub>TAF</sub> | 9:79,703-80,097 | 394.0 (35) | 79,867 (4.77) | ADGRG6 (79,770-79,923 ; 0) |
| 3 | *iHS*<sub>TAF</sub> | 9:81,292-81,834 | 542.4 (62) | 81,637 (3.20) | UTRN (81,652-82,200 ; 15) |
| 4 | *iHS*<sub>TAF</sub> | 10:67,993-68,586 | 593.6 (33) | 68,209 (3.24) | LOC112448598 (68,272-68,272 ; 62) |
| 5 | *iHS*<sub>TAF</sub> | 10:73,678-74,139 | 461.3 (56) | 73,955 (3.08) | SYT16 (73,937-74,244 ; 0) |
| 6 | *iHS*<sub>TAF</sub> | 10:76,116-76,609 | 492.9 (22) | 76,145 (2.94) | SYNE2 (76,063-76,387 ; 0) |
| 7 | *iHS*<sub>TAF</sub> | 15:34,973-35,050 | 76.90 (16) | 34,982 (3.33) | USH1C (34,963-35,012 ; 0) |
| 8 | *iHS*<sub>TAF</sub> | 16:58,363-59,261 | 897.7 (38) | 58,537 (6.27) | BRINP2 (58,417-58,560 ; 0) |
| 9 | *iHS*<sub>TAF</sub> | 23:23,433-23,866 | 432.5 (36) | 23,647 (3.51) | TRNAY-AUA (23,515-23,515 ; 132) |
|  | *Rsb*<sub>TAF/JER</sub> | 23:23,314-23,611 | 297.7 (76) | 23,555 (3.27) | TRNAY-AUA (23,515-23,515 ; 40) |
| 10 | *iHS*<sub>TAF</sub> | 23:46,236-46,627 | 390.5 (25) | 46,378 (5.15) | OFCC1 (45,984-46,226 ; 151) |
| 11 | *iHS*<sub>TAF</sub> | 23:47,230-47,618 | 387.5 (23) | 47,400 (3.61) | BLOC1S5 (47,393-47,417 ; 0) |
| 12 | *iHS*<sub>TAF</sub> | 29:27,218-28,574 | 1,355 (28) | 27,819 (2.75) | OR8B8 (27,808-27,809 ; 10) |
| 13 | *Rsb*<sub>TAF/JER</sub> | 14:49,264-49,347 | 83.10 (22) | 49,298 (2.91) | TRPS1 (48,630-48,909 ; 389) |
| 14* | *Rsb*<sub>TAF/ZMA</sub> | 6:86,541-87,032 | 491.5 (125) | 86,744 (4.19) | SLC4A4 (86,449-86,813 ; 0) |
| 15 | *Rsb*<sub>TAF/ZMA</sub> | 6:101,018-101,215 | 196.7 (54) | 101,032 (4.96) | MAPK10 (100,908-101,530 ; 0) |
| 16 | *Rsb*<sub>TAF/ZMA</sub> | 7:62,509-62,917 | 407.8 (133) | 62,534 (3.47) | SLC36A3 (62,521-62,545 ; 0) |
| 17 | *Rsb*<sub>TAF/ZMA</sub> | 8:88,183-88,415 | 232.8 (66) | 88,220 (4.34) | LOC132345997 (88,227-88,231 ; 7) |
| 18* | *Rsb*<sub>TAF/ZMA</sub> | 20:24,573-24,888 | 314.7 (75) | 24,690 (3.83) | HSPB3 (24,638-24,639 ; 51) |
| 19 | *Rsb*<sub>TAF/ZMA</sub> | 21:35,024-36,449 | 1,425 (465) | 35,244 (4.76) | STXBP6 (35,025-35,306 ; 0) |
| 20 | *Rsb*<sub>TAF/ZMA</sub> | 23:23,810-24,896 | 1,086 (360) | 23,873 (4.29) | PKHD1 (24,071-24,513 ; 198) |
| 21 | *Rsb*<sub>TAF/ZMA</sub> | 23:25,197-26,549 | 1,352 (198) | 25,333 (4.10) | GCM1 (25,311-25,333 ; 0) |

The remaining 20 candidate genes were then subjected to functional annotation using the Ingenuity Pathway Analysis (IPA) software (Ingenuity Systems, Inc., 2023). Three genes (i.e., LOC112448598, TRNAY-AUA and LOC132345997) that were not recognized in the IPA database (IPKB) were removed from the analysis. The top five diseases and biological functions significantly enriched by the analysis of the 17 annotated candidate genes (Table S9) were related to the nervous system (i.e., Neurological Disease and Nervous System Development and Function), and to tissue, organ and organism development (i.e., Organismal Injury and Abnormalities, Cell Mor-phology and Tissue Morphology). More specifically, 15 of the 17 genes (i.e., ADGRG6, AGL, BLOC1S5, GCM1, HSPB3, MAPK10, SYNE2, USH1C, UTRN, OR8B8, PKHD1, SLC36A3, SYNE2, SYT16, TRPS1) are involved in nervous system function and/or its development. Accordingly, among the top five canonical pathways identified (Table S10), four were also found to be related to the nervous system either directly (for “EGR2 and SOX10-mediated initiation of Schwann cell myelination”, Schwann cells are derived from the neural crest; “Agrin Interac-tions at Neuromuscular Junction” and “SNARE Signaling Pathway”) or indirectly (for “Interleukin-6 signaling”, IL6 is a major cytokine in the Central Nervous System).

## Discussion

The central goal of this study was to infer the demographic history and characterize the genetic makeup of the TAF cattle population that lived on Amsterdam Island until its complete and questionable eradication in 2010. Despite challenges, we were able to retrieve DNA samples for 18 individuals collected in 1992 and 2006, which we could genotype with a medium density commercial SNP assay, and for eight of them to whole-genome sequence. The sample had some limitations, and in particular we were unfortunately unable to obtain genomic information for the Y chromosome, which could have helped gain insights into the paternal lineage. However, our newly generated data allowed us to unveil intriguing details about the origins and history of the population thanks to recent method-ological advances in population genomics inference methods.

In remarkable agreement with historical records, the estimation of the TAF recent historical *N_e_* from genetic data supports a strong founder effect, down to 5 individuals, 22 generations (i.e. between 110 and 132 years for 5 to 6-year bovine generation time) before sampling, which corresponds to the late 19^th^ century. Conversely, using 1992 as the mean birth year of the analyzed individuals (Table S1) and 1871 as the founding date, such a dating can provide an indirect estimate of the cattle generation time of 5.5 years in free living conditions, which is consistent with the value classically accepted in the literature, as we discussed in Magnier *et al*. (2022). The construction of admixture graph (Patterson *et al*., 2012; Gautier *et al*., 2022) showed that two main ancestries contributed to the TAF population, consisting of about 75% to European taurine cattle (related to the present-day Jersey breed), and for the remaining to Indian Ocean zebu (related to the present-day Zebu of the islands of Madagascar and Mayotte). The timing of the corresponding admixture event (Loh *et al*., 2013) was contemporaneous with the founding bottleneck, and detailed examination of the results suggested the founders themselves may have been of heterogeneous ancestry. Such a result is consistent with the fact that Heurtin chose his animals from among those present on La Réunion island, where breeds related to European cattle (in particular to the Jersey, the “Pie-Noir bretonne” from Brittany, and the “Grise des Alpes” and Tarine both from the French Alps) and to Indian Ocean zebus were already introduced at that time (Lesel *et al*., 1969). It is further tempting to speculate that Heurtin favored individuals of Jersey or Brittany breed type, as they were best suited to survive in the harsh conditions of the Amsterdam island. The selected individuals may have had varying degrees of Indian Ocean Zebu ancestry due to more or less distant zebu ancestors, as a result of mixing on La Réunion island. Overall, the demographic inference of the genetic history of the TAF population confirmed that this population represents a rare example and outstanding case study of a successfully established large mammal from a recent, brief and extreme population bottleneck that has rapidly thrived in a seemingly challenging environment. In the recent population genomics literature dealing with mammalian populations, either invasive or in conservation, that have experienced highly dynamic demographic histories, the age of the bottleneck (22 generations), its intensity (only 5 founders), its short duration (only a few generations) and the fact that the population recovered make the TAF example unique.

We further performed a detailed comparative analysis of genetic diversity at the whole-genome level of the TAF population based on the WGS data of the 8 TAF individuals. Despite high levels of individual inbreeding (approx-imately 30%), the TAF population showed no evidence of strong diversity reduction. Autozygosity was associated with HBD segments of approximately 3 Mb in length, consistent with the recent founder event. Also, and in agree-ment with a rapid expansion scenario, we found little evidence of recent inbreeding (i.e., absence of long HBD segments), suggesting that no new inbreeding occurred a few generations after the bottleneck and levels stabilized. This was confirmed by comparing the distribution of HBD segments in individuals sampled at different times. This also illustrates that the bottleneck-expansion scenario leaves a characteristic footprint in terms of the distribution of HBD segments (i.e. concentrated in one HBD class), as observed for example in the Soay sheep population (Druet and Gautier, 2017), which is well captured by the multiple HBD class HMM. The observed levels of inbreeding are similar to those reported in endangered or previously endangered populations such as mountain gorillas (Xue *et al*., 2015), Royal Isle wolves (Robinson *et al*., 2019), European bison (Druet *et al*., 2020) and Royal Isle moose (Kyriazis *et al*., 2023), and higher than for Alpine ibex (about 15%) (Grossen *et al*., 2020), vaquita (5%) (Robinson *et al*., 2022) and Ethiopian wolves (16%) (Mooney *et al*., 2023). However, the TAF population showed relatively high genetic diversity compared to populations with similar levels of inbreeding. Indeed, the median individual heterozygosity of 0.2% was more than an order of magnitude higher than that of the Royal Isle moose, Alpine ibex, vaquita, Ethiopian wolf, Iberian lynx (Kleinman-Ruiz *et al*., 2022) and island fox (Robinson *et al*., 2016), which have reported heterozygosity levels < 0.05%. Such levels of heterozygosity (> 0.1%) were only observed in the European bison (Gautier *et al*., 2016a) or the wolves of the Royal Isle (Robinson *et al*., 2016). The TAF population thus has both relatively high genetic diversity and inbreeding levels, two parameters that are usually inversely related, resulting in a unique combination of inbreeding and heterozygosity levels.

The limited reduction in genetic diversity is consistent with the inferred admixed origins of the founders and the subsequent rapid population growth. Indeed, reductions in heterozygosity have been most commonly observed in populations that have experienced reductions in effective population size maintained over many generations, as in the examples above. In agreement, Robinson *et al*. (2023) predicted that bottleneck of short duration minimally impact genetic diversity. In addition to high levels of inbreeding associated with moderate diversity reduction, we observed that the SFS was strongly flattened. The TAF population has proportionally fewer singletons and more variants segregating at moderate frequencies than the other populations we analyzed. This has been reported in other populations that experienced a bottleneck, such as the Ethiopian and Royal Isle wolves (Mooney *et al*., 2023)and the Australian rabbit (Alves *et al*., 2022). The flattening of the SFS curve is a signature of strong drift, which could occur despite only a few generations of reduced *N_e_*. A final clear signature of the strong founder event on the genetic diversity of the TAF population is the high level of fixation. Indeed, we estimated that four times more derived alleles reached fixation in TAF than in the two zebu populations studied, but this level was still below that observed in the two European cattle populations.

We observed that the SFS curve was flattened in the TAF population for all classes of variants, including delete-rious NS and LoF variants. Compared to the European taurine (JER and HOL) or Indian Ocean zebu (MAY and ZMA) control populations, where, as expected, the proportion of singletons in these two classes was higher (rela-tive to intergenic variants), the enrichment was less pronounced in the TAF sample. This drift may be accompanied by a relaxation of the purifying selection on deleterious variants. Accordingly, the proportions of segregating NS variants (a proxy for mildly deleterious variants) and the NS/S ratio were found to be higher in the TAF popu-lation compared to the two zebu populations, as indicated by the *R_X_*_/*Y*_ statistic which showed a slight excess of NS variants. Interestingly, these statistics were less extreme in the TAF than in the European breeds JER and HOL, since these dairy cattle have been subjected to intense selection for several decades and have maintained a small *N_e_* of only a few tens to hundreds of individuals over the last tens of generations (Gautier *et al*., 2007). Relaxation of selection on NS variants has previously been observed in domesticated species as a result of *N_e_* reduction (Cruz *et al*., 2008; Marsden *et al*., 2016; Bosse *et al*., 2019), and has also been reported, for example, in mountain gorilla (Xue *et al*., 2015). However, evidence of purging of the most deleterious variants (e.g. LoF variants), with *R_X_*_/*Y*_ ≪ 1 were observed in other populations with small *N_e_* such as mountain gorilla (Xue *et al*., 2015), Ibex (Grossen *et al*., 2020) and kākāpō (Dussex *et al*., 2021). Such purging has also been found in other endangered populations, such as the Iberian lynx (Kleinman-Ruiz *et al*., 2022), the Royal Isle moose (Mooney *et al*., 2023) and the vaquita (Robinson *et al*., 2022). Yet, despite the high level of inbreeding in TAF, the purging of LoF variants appears limited. Indeed, there were no strong signal of reduction of these highly deleterious variants. Although lower proportions of segregating LoF variants and *R_X_*_/*Y*_ statistics slightly below 1 suggest mild purging, the higher proportion of fixed LoF variants and the SFS curves suggest rather a relaxation of purifying selection. These two processes, drift and negative selection, independently affect the outcome of deleterious variants (Grossen *et al*., 2020), alongside the pivotal role of “chance”, which makes the consequences of a bottleneck highly variable (Bouzat, 2010; Robinson *et al*., 2023). Overall, the absence of a clear signal of purging is consistent with the fact that this requires a reduction in *N_e_* lasting many generations (e.g. Bouzat, 2010; Mooney *et al*., 2023; Robinson *et al*., 2023; Smeds and Ellegren, 2023), whereas in the case of the TAF population, the bottleneck may have been too short for purging to be effective. Indeed, slower rates of inbreeding enhance the effectiveness of purging and longer periods of reduced *N_e_* offer more opportunity for purging the genetic load (Bouzat, 2010). Similar observations (i.e., absence of clear signal of purge and high fixation levels) have recently been reported in the Scandinavian wolf that experienced a very recent reduction in population size (Smeds and Ellegren, 2023). It is interesting to note that despite the absence of clear signal of purge, the relaxation of selection and the resulting substantial realized load that we observed, the population was still able to expand and recover after several population declines.

If the extreme bottleneck associated with the arrival of cattle on the island of Amsterdam did not pose an insurmountable demographic challenge to this population, it still had to face and adapt to extreme environmental conditions. Indeed, the few founding individuals may have benefited from the farmer’s care for only a few months before being left alone. The remote location of the sub-antarctic island of Amsterdam in the Southern Ocean exposes it to harsh and unpredictable weather patterns. The island is frequently buffeted by strong winds, sometimes reaching hurricane force, and the persistent cold (with temperatures often hovering around freezing) adds to the challenging conditions. In addition, the island’s isolation and limited resources, particularly fresh water, add to the harshness of the environment. Nevertheless, the estimation of domestic bovine maladaptation in the form of Genetic Offset (Capblancq *et al*., 2020), showed it was among the lowest (i.e., less challenging) for the environmental conditions associated with the geographical origin (i.e., Channel islands and north of Brittany) of the main European taurine ancestry of the TAF population. It should be noted that we have computed GO here assuming that the sample of bovine populations analyzed, representative of worldwide diversity but with a deliberate focus on EUT breeds, are adapted to the environment of their birthplace of origin (in the sense that adaptive alleles are at their local optimum frequency), while also including the TAF populations to allow the representation of the Amsterdam island conditions. This is reasonable since the primary goal of the GO computation is to model the relationship between environmental conditions (characterized by a set of 19 bioclimatic covariates each averaged over the period 1981-2010) and the structuring of genetic diversity in the underlying populations via a GEA model. This allows environmental distances to be appropriately weighted to account for genetic adaptation in the derivation of GO (Gain *et al*., 2023). Overall, although the absolute value of GO is difficult to interpret per se, it has been shown to be directly related to average population fitness (Gain *et al*., 2023) and to the establishment probability of invasive species (Camus et al., in prep.). Taken together, the GO analyses suggest that the adaptive challenge on Amsterdam island was limited given the inferred origin of the TAF population. In other words, the successful establishment of the TAF population can potentially be explained by a form of pre-adaptation of its founders to the local climatic conditions due to their predominant European taurine ancestry. The TAF population thus provides a good example of the pre-adaptation of a non-indigenous population able to survive and reproduce in local environmental conditions close to those of its place of origin. This is consistent with the so-called matching hypothesis, despite low propagule pressure (Sol, 2007). In addition, and in agreement with Bouzat (2010), a small adaptive challenge might also explain lower levels of inbreeding depression.

Recent studies have highlighted the “island syndrome”, which includes the frequent dwarfism of large-bodied mammals on islands (Rozzi *et al*., 2023). Since the TAF is a rather short cattle population, it was tempting to speculate that the size resulted from a rapid dwarfism (Rozzi and Lomolino, 2017) and that the feralization process and food restriction in the island may indeed have favored selection for small animals. However, our results argue against such an insular dwarfism syndrome in the TAF population. First, the demographic inference strongly suggests that the small size of TAF individuals, with a wither height of 134 cm and 113 cm (Lesel *et al*., 1969) and an average weight of 389.6 kg±42.8 and 293 kg ±45.8 (Berteaux and Micol, 1992) for males and females, could more parsimoniously be directly related to the small format of the populations from which they originate. The JER breed is among the smallest of all dairy breeds with an average female wither height of around 120 cm and female weight of 375 kg. In addition, old but detailed reports confirm that the most numerous cattle population that lived in Brittany (to which the current BPN is related) at the end of the 19^th^ was notoriously small, with a reported average size of 100 to 110 cm on average (de Lapparent, 1902). Similarly, Indian Ocean zebus are of small format with, for example, an average male wither height of 110 cm and weight of 240 kb measured in the ZMA (Zafindrajaona and Lauvergne, 1993). Second, at the genome level, our estimates of breeding values for stature based on estimated effects for 105 variants obtained in an extended meta-analysis that included several cattle breeds (Bouwman *et al*., 2018) did not provide evidence for strong (polygenic) selection for short stature.

Although the environmental conditions may not have been as challenging, the living conditions of the TAF popu-lation changed dramatically and rapidly. This led to a complete feralization of the initially domesticated animals in just a few generations, making this population of paramount interest for yet another aspect. To complete the genomic characterization of the population, we therefore examined the genomic response of the TAF population to the new adaptive constraints it encountered. Strikingly, the majority of candidate genes we were able to identify within the footprints of selection were annotated to be involved in nervous system function and development. We can view these results as consistent with the brain size–environmental change hypothesis, which states that large brains in mammals are associated with an enhanced behavioral flexibility, which may confer advantages to indi-viduals by improving their fitness in the face of novel environmental conditions (Sol *et al*., 2008). They are also consistent with the behavioral modification of TAF individuals that may have accompanied and contributed to the rise of the population on the island and its feralization. Several observers have noted a clear and complex social organization of the TAF, similar to that of wild bovids, with groups of females in matrilineally structured groups consisting mostly of females and young to subadult males; geographically separated groups consisting exclusively of adult and/or subadult males; and male-female groups usually formed at the beginning of the reproductive season by incorporating adult males into the groups of females (Daycard, 1990). Such a complex social structuring, that accompanied feralization, is also supported by the estimated negative population *F_IS_* (Chikhi and Parreira, 2015) and the close relatedness of some samples, although our sample size remained limited. Observers also reported that the TAF individuals had become fierce and phenotypically, there was a clear and impressive diversification of color patterns in all the individuals (Lesel *et al*., 1969). If feralization cannot be viewed as a mere reversal of domestication (Gering *et al*., 2019), these observations and our results showing the apparent importance of nervous system function in TAF adaptation reveal common features between the feralization process in the TAF population (fierceness) and the domestication syndrome observed in domesticated mammals (tameness), mobilizing genes involved in the neural crest development (Wilkins *et al*., 2014). They also highlight the likely polygenic nature of complex traits involved in feralization, and the role of standing variation in the rapid adaptation of the TAF population to the wild. Indeed, this may explain how the variants underlying a feralization that lasted only a few generations were still segregating in the domestic cattle from which the founders originated.

The TAF population represents an unprecedented resource of a domestic population that was able to colonize and thrive in a challenging environment, recovering from only a handful of domestic founders without strong evidence for purging. The population quickly became feral, developing new abilities rendering it adapted to an harsh environment. Unfortunately, despite a convincing and conclusive management policy to allow cohabitation with endemic species, this population was eradicated in 2010. Moreover, no effort was made to conserve some individuals or even, at the very least, to keep samples indicating that those who promoted this eradication, including biologists, considered it a nuisance without any scientific interest. However, we hope that this article will convince the reader of the opposite and contribute to a more careful reflection before eradicating feral populations. The data and analyses we present will help to preserve a trace and a legacy of the incomparable resource that this population represented for the scientific community.

## Materials and Methods

### Genetic data

#### New sample origin, genotyping and sequencing data

Blood samples for 18 different bovine individuals from the Amsterdam island (TAF) were collected during two sampling campaigns in 1992 (n=12 females) and 2006 (n=6, 3 females and 3 males) and frozen at-20°C before being transferred to the former LGbC laboratory (INRA, Jouy-en-Josas) and to the LaboGENA genotyping platform (Jouy-en-Josas, France), respectively (Table S1). Genomic DNA for the 12 females collected in 1992 were extracted in March 1994 using the protocol described in Jeanpierre (1987) which allowed long-time preservation of DNA integrity at-20°C. The genomic DNA of the 6 individuals collected in 2006 was extracted at the LaboGENA platform for genotyping purposes and was unfortunately neither stored nor returned after the test. All the 18 TAF individuals were genotyped on the Illumina BovineSNP50 chip assay v2 (Matukumalli *et al*., 2009) at the LaboGENA platform (Jouy-en-Josas, France) using manufacturer recommendations (Illumina, 2016), in 2014 for the 12 individuals collected in 1992 and in 2010 for the 6 individuals collected in 2006. Together with this latter TAF individuals, 31 individuals belonging to the Moka Zebu breed (MOK) sampled in 2010 in La Réunion island were also genotyped on the same assay. All these newly generated genotyping data (n=49 individuals in total) have been made publicly available in the WIDDE repository (Sempéré *et al*., 2015). Finally, 8 of the 12 TAF females collected in 1992 and for which enough DNA was still available were further paired-end sequenced (2×150 nt) on a HiSeqX Illumina sequencer at the Macrogen commercial platform (Macrogen Inc., Seoul, South Korea). For the purpose of this study, we also sequenced sixteen zebus individuals from Mayotte (n=8) and Madagascar (n=8) is-lands that were collected in 2017 and 1989 respectively, and that were among the individuals previously described and genotyped in Magnier *et al*. (2022) and Gautier *et al*. (2009). Thirteen (8 MAY and 5 ZMA) were paired-end sequenced on a HiSeq 2500 (n=5 with 2×125 nt) or a NovaSeq 6000 (n=8 with 2×150 nt) Illumina sequencer at the MGX platform (Montpellier, France). The three remaining ZMA individuals were paired-end sequenced (2×150 nt) on a NovaSeq 6000 Illumina sequencer at the Genoscope platform (Evry, France) (Tables S3 and 1). All the newly generated sequencing data for the 8 TAF, the 8 MAY and the 8 ZMA individuals were deposited in the NCBI Sequence Read Archive repository under the BioProject accession number PRJNA1010533 (Table S3).

#### The 50K SNP genotyping dataset (W50K) representative of worldwide cattle genetic diversity

To explore the genetic relationship of the TAF population with other bovine population and infer its origin, the newly generated data obtained with the BovineSNP50 assay were combined with publicly available data from populations representating the worldwide cattle diversity (Gautier *et al*., 2010; Matukumalli *et al*., 2009) and that are stored in the WIDDE database (Sempéré *et al*., 2015). As detailed in Table 1 and represented in Figure 3A, the resulting W50K data set finally consists of 876 individuals from 32 different populations. Genotyping data were filtered using WIDDE utilities (Sempéré *et al*., 2015), only retaining individuals with a SNP genotyping call rate > 95%. Likewise, we only retained SNPs with an individual genotyping call rate > 75% in all population samples (leading to an overall genotyping call rate > 90%) and that mapped to the latest ARS-UCD1.2 (aka *bosTau9*) bovine genome assembly (Rosen *et al*., 2020). In addition, we discarded SNPs with a Minor Allele Frequency (MAF) < 0.1% or showing a highly significant departure of individual genotype frequencies from Hardy-Weinberg equilibrium expectations (p< 10^−4^) in at least one population. The W50K data set finally comprised 40,484 SNPs including 40,426 autosomal SNPs (from 736 on chr. 8 to 2,628 on chr. 1).

#### Whole genome sequencing data processing, variant calling and annotation

To provide a comprehensive genome-wide analysis of TAF genetic diversity and compare it with the most closely related populations as inferred from population genetics analyses on the W50K genotyping data, we analyzed the newly generated WGS data for the 8 TAF, 8 ZMA and 8 MAY together with 8 Jersey and 8 Holstein publicly available WGS data (Daetwyler *et al*., 2014) that displayed similar coverage (Tables S3). To infer the ancestral state of the identified variable position, we used WGS data for one American bison (*Bison bison*), one European bison (*Bison bonasus*), one Gaur (*Bos gaurus*), and one Banteng (*Bos javanicus*) availble from (Wu *et al*., 2019). Processing of all the WGS data mostly followed the “1000 Bull Genomes” analysis guidelines (version of 2018/06/18) (Daetwyler *et al*., 2014). Briefly, adaptors were removed from the sequencing reads that were trimmed and filtered using Trimmomatic v0.38 (Bolger *et al*., 2014) that was run with options LEADING:20, TRAILING:20, SLIDINGWINDOW:3:15, AVGQUAL:20, MINLEN:35 and ILLUMINACLIP with the TruSeq3-PE.fa:2:30:3:1 adaptor file. The program fastp v0.19.4 (Chen *et al*., 2018) was further run with default options on the resulting fastq files mainly to provide statistics for Quality Check and also to (marginally) improve adapter removal. After filtering, paired reads were aligned onto the ARS-UCD1.2_Btau5.0.1Y whole genome assembly that combined the ARS-UCD1.2 (for autosomes and the X chromosome) (Rosen *et al*., 2020) and the Btau5.0.1 Y chromosome assemblies (Bellott *et al*., 2014), using the program bwa mem (Li, 2013) run with default options. PCR and optical duplicates were subsequently marked using the command MarkDuplicates from Picard 2.18.2 (Broad Institute, 2018). Base quality scores were further recalibrated (BQSR step) with the BaseRecalibrator tool from GATK (v4.2.6.1) (McKenna *et al*., 2010) using a catalogue of 110,270,189 known variants from the 1000 Bull genome project (Daetwyler *et al*., 2014) as a recalibration file named ARS1.2PlusY_BQSR.vcf. After BQSR, we performed variant calling using GATK’s HaplotypeCaller setting the ploidy to one for the mitochondria and for the X chromosome in males. A multi-sample vcf file was then generated for each chromosome (and the mitochondria) by combining all the 40 resulting gvcf files for bovine individuals with the GATK’s CombineGVCFs and GenotypeGVCFs tools.

Finally, mitochondrial haplogroups from each individual were inferred using MitoToolPy (Peng *et al*., 2015) based on the “treeFile” provided by Dorji *et al*. (2022) for the ARS_UCD1.2 assembly. For each individual, a fasta file was obtained by extracting genotypes from the VCF file using the consensus command from bcftools v1.10.2 and using the reference genome fasta file.

### Population genetics structure

#### F-statistics computation

The Wright fixation indexes *F_IT_*, *F_S T_* and *F_IS_* were estimated using a custom implementation of the estimator proposed by Weir and Cockerham (1984) under an analysis of variance framework (see also Weir, 1996, p176– 179) between all pairs or over all the samples. Within-population *F_IS_* were estimated following Weir (1996, p.80) and heterozygosities were estimated with the compute.fstats function of the R package poolfstat (v2.2.0) (Gautier *et al*., 2022). Standard errors (s.e.) and 95% confidence interval (as ±1.96 s.e.) of the different statistics values were estimated using a block-jackknife approach (Busing *et al*., 1999; Gautier *et al*., 2022). This here consisted of dividing the genome into contiguous chunks of 250 SNPs, leading to 150 blocks of 15.3 Mb on average (from 10.6 to 21.4 Mb), and then removing each block in turn to quantify the variability of the estimator among the 150 corresponding estimates.

#### Exploratory analyses

The neighbor-joining trees (Saitou and Nei, 1987) were computed using the nj function of R package ape (Par-adis *et al*., 2004) based on the matrix of allele sharing distances (ASD) between all pairs of individuals following (Gautier *et al*., 2010). Principal component analysis (PCA) of individual SNP genotyping data was carried out as described in Patterson *et al*. (2006) using the svd function of the R package base (R Core Team, 2017). To provide an alternative description of the structuring of genetic diversity, unsupervised genotype-based hierarchical clustering of the individuals was carried out using the maximum-likelihood method implemented in the ADMIXTURE (v1.06) software (Alexander *et al*., 2009). Results were visualized with custom functions in the R environment (R Core Team, 2017).

#### Relationship inference

The program King (v2.3.2) (Manichaikul *et al*., 2010) was used to infer relationship among pairs of TAF individuals based on autosomal genotyping data. Inference relied on both the estimates of kinship coefficient and IBD-segment sharing (--kinship--ibdseg options respectively). We also considered the option--unrelated to define a maximal set of unrelated individual (up to third degree relationships).

### Demographic inference

#### *f*-statistics-based tests and admixture graph construction

*f*-statistics based demographic inference (Patterson *et al*., 2012) were carried out with the R package poolfstat (v2.2.0) (Gautier *et al*., 2022). We used the compute.fstats function to estimate the different *f*-statistics including *F*_3_ for all the population triplets and *F*_4_ for all population quadruplets. As for the Wright fixation indexes previously described, standard-errors of the estimated statistics (and their corresponding Z-scores for *f*_3_) were estimated using block-jackknife defining blocks of 250 consecutive SNPs (i.e., option nsnp.per.bjack.block=250). Following Patterson *et al*. (2012), formal tests of population admixture were carried out using the estimated *f*_3_ statistics, a negative Z-score (*Z* < −1.65 at the 95% significance threshold) associated to an *f*_3_ for a given population triplet A;B,C indicating that the target population A is admixed between two source populations each related to B and C. Admixture graph construction and exploration were carried out with poolfstat utilities using a semi-automatic approach similar to that described in Gautier *et al*. (2022). We relied in particular on the graph.builder function (ran with default options) to position populations onto scaffold graphs. The fit of the best fitting graphs (based on the BIC criterion) was further validated with the *compare.fitted.fstats* function to compare to which extent the estimated *f*-statistics depart from their predicted values based on the fitted admixture graph parameters via a Z-score (Patterson *et al*., 2012; Lipson, 2020; Gautier *et al*., 2022). The main steps of the graph construction are illustrated and detailed in Figure S5. Based on the corresponding inferred history and as detailed in Figure S6, the EUT populations represented in the W50K were ranked for their proximity with the European ancestral source of the TAF or MOK using *f*_3_ and *f*_4_ estimates for all (X;ZMA,Y) and (Y,NDA;X,GIR) configurations respectively (where X=TAF or X=MOK and Y is the tested EUT population). Finally, the 95% CI of the proportion of EUT ancestry was (re)estimated more accurately for TAF (and MOK) using *F*_4_-ratios as described Figure S6 with the compute.f4ratio poolfstat function (Patterson *et al*., 2012; Gautier *et al*., 2022).

#### Estimation of the timing of admixture

We estimated the timing of admixture events (in generations) with the program alder v(1.03) (Loh *et al*., 2013). This approach relies on the modelling of the exponential decay of admixture-induced LD in a target admixed population as a function of genetic distance, using a LD measure weighted by allele frequencies in either one or a pair of source population proxies. Genetic distances between pairs of SNPs were derived from physical distances assuming a cM to Mb ratio of 1 and a six-year generation time was assumed to convert the timing from generations to years (Magnier *et al*., 2022).

#### Inference of the recent population size history

Historical effective population sizes (*N_e_*) were inferred with the program Gone that implements the approach developed by Santiago *et al*. (2020) to fit the observed spectrum of LD of pairs of loci over a wide range of recombination rates (which we derived from physical map distances assuming a cM to Mb ratio equal to 1 as above). Following Magnier *et al*. (2022), we adopted a block-jackknife approach to estimate confidence intervals for the inferred *N_e_* trajectories by defining non-overlapping blocks of 250 consecutive SNPs as above (block size of ca. 15 Mb).

#### Age-based partitioning of individual inbreeding

To identify and classify homozygous-by-descent (HBD) segments, we relied on the model-based approach that is implemented in the R package RZooRoH (v0.3.2.1) (Druet and Gautier, 2017, 2022; Bertrand *et al*., 2019). Within individual genomes, HBD segments correspond to segments inherited twice from a common ancestor as a result of inbreeding, and are often detected as runs-of-homozygosity (ROH), which are used as a proxy for HBD. In the ZooRoH model, autozygosity is partitioned into multiple HBD classes. HBD classes are defined by their rate parameter *R_c_*, which determines the expected length of HBD segments, and correspond to groups of ancestors present in distinct past generations. To do this, we converted the VCF into a genotype probabilities file (GEN format) using bcftools (v1.10.2) and the Phred-Likelihood (PL) field, retaining only the 1,091,824 SNPs included in commercial genotyping arrays (Nicolazzi *et al*., 2014) and fitted the “layer” model (Druet and Gautier, 2022) specifying 13 HBD classes with rates *R_c_* equal to {2, 4, 8,..., 8192}.

### Estimation of genetic diversity from WGS data

Genetic heterozygosities based on WGS data were estimated from each individual genome alignments (bam) files described above that were further filtered for mapping quality and duplicate reads with the program view (run with option-q 20) and rmdup of the samtools (v1.13) suite (Li *et al*., 2009). To that end we relied on the program mlrho version 2.8 (Haubold *et al*., 2010) that implements a maximum likelihood estimator of the population mutation rate (θ = 4*N_e_*µ) which fairly approximates heterozygosity under an infinite sites model (and providing θ is small), while simultaneously estimating sequencing error rates and accounting for binomial sampling of parental alleles (Lynch, 2008). Following Gautier *et al*. (2016b), only sites covered by 3–30 reads (after discarding bases with a base quality BAQ< 25) were retained in the computation.

For further analyses of genetic diversity and genetic load, the VCF was recalibrated using the GATK’s VariantRecalibrator command (VQSR step) using the file that was used for BQSR as known set and a file with 1,213,314 SNPs from commercial arrays as the truth set (Nicolazzi *et al*., 2014). We then selected bi-allelic SNPs with a quality score corresponding to 99% conservation of the truth set and that mapped to autosomes. In addition, SNPs called for less than 90% of individuals or with an average individual depth of coverage (DP) < 5 or ≥ 20 were discarded, leaving 23,383,523 SNPs for further analyses.

Allele frequencies (AF) were estimated per population from individual genotype likelihoods (PL) using a custom implementation (in modern Fortran) of the EM-algorithm described by Kim *et al*. (2011). The number of segre-gating sites per population was estimated as the number of sites with a 0.001 < *AF* < 0.999. The number of fixed sites per population was derived from the proportion of sites with a frequency of the derived allele greater than 0.999. For allele polarisation, we repeated the variant calling by adding the BAM files of one individual from four outgroups (*Bison bison*, *Bison bonasus*, *Bos javanicus* and *Bos gaurus*) (Table 1). When all outgroup individuals were homozygous for an identical allele, and at most one outgroup had a missing genotype, we called this allele the ancestral allele. Using this approach we were able to define the ancestral allele for 83.8% of variants. The number of fixed sites per population was then estimated as the proportion of fixed derived alleles multiplied by the total number of sites. To compare the Site Frequency Spectrum (SFS), we calculated the probability of the presence of 1 to 15 derived alleles in each population in order to have only discrete values. This was therefore applied only to variants for which the derived allele was identified and without missing genotypes in that population. These genotype probabilities were derived from the phred likelihoods (PL) of the eight individuals from the population and by enumerating the 6561 possible genotype combinations (3 possible genotypes for 8 individuals). For each combination, the number of derived alleles is known and the probability can be estimated as the product of the eight individual vector of genotype probabilities (GP). Finally, the probability of the presence of N derived alleles in the population was estimated as the sum of probabilities of all combinations with N derived alleles.

### Genetic load and distribution of deleterious variants

We relied on the Variant Effect Predictor (VEP) v95.0 (McLaren *et al*., 2016) to annotate our variants. Variants annotated as “stop gained”, “splice donor” or “splice acceptor” were defined as Loss-of-Function (LoF) variants. Additional classes consisted of “intergenic”, “synonymous” (S)and “non-synonymous” (NS) variants, which could be further subdivided into “deleterious non-synonymous” and “tolerated non-synonymous” classes. In the case of multiple annotations, we retained the most severe annotation. This classification of variants was first used to compare, across the different populations, the proportions of segregating and fixed variants in different classes and the SFS obtained for each class. We then calculated measures related to the relaxation of purifying selection, the occurrence of purging of deleterious alleles and to the masked and realized loads. First, we estimated the non-synonymous to synonymous ratio (NS/S) as the ratio of heterozygous genotypes for these two categories per individual (Cruz *et al*., 2008; Renaut and Rieseberg, 2015). Next, the masked and realized load, measured for a group of deleterious variants, were estimated as in Smeds and Ellegren (2023). The masked load of an individual is defined as the proportion of heterozygotes genotypes whereas the realized load is the proportion of genotypes homozygous for the derived allele (both statistics were calculated using only the called genotypes of that category within each individual). These first measures using genotype counts per individual were calculated using genotype probabilities. Finally, we calculated the relative number of derived alleles *R_X_*_/*Y*_ (Do *et al*., 2015) in two populations *X* and *Y*. To do this, the probability of sampling a derived allele in population *X* and not in population *Y* is calculated for each SNP using the estimated AF of the derived alleles in population *X* ( *f_X_*) and *Y* (*f_Y_*) as (*f_X_*(1 − *f_Y_*)) and then summed over all SNPs. The opposite probabilities, ( *f_Y_* (1 − *f_X_*)), are summed over all SNPs and *R_X_*_/*Y*_ is finally obtained as the ratio of these two values. We estimated *R_X_*_/*Y*_ for missense and LoF variants, and these values were standardized by the values obtained for intergenic variants (as proxy for neutral variants). This standardization corrects for differences in branch lengths (Do *et al*., 2015). Note that we obtained *R_X_*_/*Y*_ values close to 1 for intergenic variants, indicating that standardization was not necessary. The confidence interval of the *R_X_*_/*Y*_ values was obtained by a block jackknife procedure as in Xue *et al*. (2015). Briefly, we divided the genome into 100 blocks of consecutive SNPs, with the same number of SNPs per block, and repeated the calculation of *R_X_*_/*Y*_ by ignoring each block one by one.

### Adaptive history

#### Estimation of genetic o**ff**set

The relative degree of “maladaption” to the environment of Amsterdam island was evaluated using a genetic offset (GO) statistic (Gain *et al*., 2023) for all the cattle populations represented in the W50K dataset. This statistic quantifies the difference between a source and a target environment as a weighted distance between their underlying environmental (e.g., bioclimatic) covariable values. The weights on the covariables are directly related to the importance of their association with the (adaptive) genomic composition of the breeds that is estimated under a GEA (Gene-Environment Association) model assuming the different populations are adapted to their environment of origin. Following Gain *et al*. (2023), we relied on a linear modelling of the relationship between the genetic diversity of the 32 cattle population in the W50K dataset and 19 bioclimatic covariables (averaged values over the period 1981-2010 at a 30 arc sec resolution) that were extracted from the CHELSA (v2.1, accessed the 21st May 2023) database (Karger *et al*., 2017, 2018) to characterize their environment based on the GPS coordinates of their birthplace (Table 1). More precisely, we first carried out a PCA on the scaled and centered covariables using the R package ade4 (Dray and Dufour, 2007), and retained the first seven PCs that together explained 98.7% of the overall variation. We then fitted a linear model using the covmcmc model implemented in BayPass v2.4 (Gautier, 2015) and used the resulting (posterior mean) estimates of the vectors ***β_k_*** of the *n_snp_* = 40, 426 SNP regression coefficients for each of the seven (scaled) PCs *k*. In short, the regression coefficients summarize the effect of each environmental covariable on the distribution of allele frequencies corrected for the neutral structuring of genetic diversity. The GO for each population *j* environment with respect to the Amsterdam island environment characterized by the vectors ***e_j_*** and ***e_ta f_*** of seven PCs, respectively, was then computed as

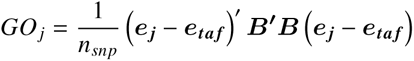

where ***B*** is the (*n_snp_* × 7) matrix with all the seven ***β_k_*** vectors stacked side by side (Gain *et al*., 2023). As a matter of comparison we also computed the (unweighted) environmental distance as 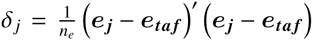 that simply corresponds to the Euclidean distance between environmental PCs (where *n_e_* = 7 is the number of environmental PCs). To compute the GO distribution over the whole European continent, we extracted the 19 environmental bioclimatic covariates for all positions across the entire European continent from CHELSA v2.1. These were then transformed into PCs using the above PCA loadings (with the suprow function of ade4) that we rescaled as the breed bioclimatic PCs to compute GO. The resulting GO rasters were processed and analyzed at a resolution of 0.01°with utilities from the raster R package (Hijmans, 2022).

#### Genomic prediction of breeding values for height based on WGS data

To investigate whether the TAF are enriched in alleles associated with short stature as a result of possible selection, we compared the frequency of such alleles in 8 sequenced individuals with the frequency observed in the four other breed samples from the WGS data set. To do this, we used a list of 164 variants identified in a large cattle meta-analysis (Bouwman *et al*., 2018) (mapped to the UMD3.1 bovine reference genome). Among them, 105 variants, with matching reference alleles in both reference genomes, were found polymorphic in our data. Individual genotypes for these alleles were then used to predict height differences between individuals, as the sum over all SNPs of the allele dosage by the reported allele effect on height. This is equivalent to a comparison allele frequencies, weighted by their effect as precisely estimated by (Bouwman *et al*., 2018) on various cattle breeds.

#### Genome scan with EHH-based test using 530K SNP haplotypes

To increase the power and sensitivity of the detection of footprints of selection, we first built a data set consisting of 71 individuals (18 TAF, 23 ZMA and 30 JER) that were genotyped, either directly or via imputation, for 530,769 autosomal SNPs from the Illumina BovineHD SNP assay (comprising >770,000 SNPs). To that end, we first extracted genotypes at 718,718 BovineHD SNPs that we could unambiguously identify in the vcf file we generated with our WGS data and recoded them in the so-called “Top” format of the chip. We then combined the obtained genotypes for the 8 TAF, 8 ZMA, 8 MAY and 8 JER sequenced individuals with SNP genotyping data for the 18 TAF on the BovineSNP50K assay and for the 30 JER, 32 MAY and 26 ZMA individuals on the BovineHD assay that were publicly available from previous studies (Magnier *et al*., 2022) and were extracted from the Widde database (Sempéré *et al*., 2015). We used genotypes from 24 the individuals (8 TAF, 8 MAY and 8 ZMA) that were both sequenced and genotyped to assess the genotype concordance rate. Among their 8,843,201 genotypes available from both WGS data (with a GQ > 20) and the genotyping array, only 0.42% were found to be different. For those, we chose to keep the SNP assay genotype in the combined data set further excluding five individuals (3 ZMA and 2 MAY) that remained poorly genotyped. Likewise, we only retained the 547,214 SNPs that were genotyped on more than 90% of the individuals in each of the four population samples.We then proceeded with the imputation of missing genotypes for the 10 TAF individuals that were only genotyped on the 50K SNP assay (corresponding to the targets). The reference panel consisted of the the 40 JER, the 30 MAY, the 8 TAF and the 23 ZMA individuals genotyped for the 530,769 SNPs. We first carried out haplotype phasing of the 29 bovine autosomes with Beagle v5.4 (Browning *et al*., 2021) for the reference and target panels separately. We then carried out missing genotypes imputation using Beagle v5.4 (Browning *et al*., 2018).

Based on the resulting haplotypes, we then carried out genome-wide scan for footprints of positive selection using EHH-based tests within the TAF population using the *iHS* (Voight *et al*., 2006) statistic for within popula-tion and the *Rsb* statistic (Tang *et al*., 2007) for pairwise-population analyses, as implemented in the R package rehh v3.1.2 (Gautier *et al*., 2017; Klassmann and Gautier, 2022). For the standardization of *iHS*, we only retained the 204,867 SNPs with a MAF>0.01 and the alleles were not polarized (i.e. scan_hh was run with polarized=FALSE and ihh2ihs was run with options min_maf=0.01 and freqbin=1) as discussed in Klass-mann and Gautier (2022) and section 7.6 of the online rehh vignette. The *iHS* statistics was further transformed into p_iHS_ = − log_10_ (1 − 2 | Φ (iHS) − 0.5 |) (where Φ(*x*) is the Gaussian cumulative function). Assuming *iHS* is normally distributed under neutrality, the resulting p_iHS_ = − log_10_ (2 | Φ (− | iHS |)) may then be interpreted as two-sided a P-value (on a negative log_10_ scale) associated with the neutral hypothesis of no selection. For the analyses based on *Rsb* derived form the ratio of TAF integrated site EHH in the numerator and either that of JER or ZMA in the denominator, we instead focused on one-sided P-value of the form p_Rsb_ = − log_10_ (1 − Φ (Rsb)) to identify TAF-specific signals only. Candidate regions were identified by combining the obtained SNP-specific P-values into a local-score derived from a Lindley process as described in Fariello *et al*. (2017) with a SNP score equal to − log_10_(*p*) − *ξ*. We here chose *ξ* = 2 and only retained the windows with a local-score peak value significant at a 1% P-value threshold (analytically derived for each chromosome separately). The obtained windows were then annotated for their number of SNPs, their size, the position of the *iHS* or *Rsb* peak, and the closest gene using the NCBI annotation gff file for the ARS-UCD1.2 genome assembly.

The candidate genes were functionally annotated using Ingenuity Pathway Analysis software (Ingenuity Systems, Inc., 2023) considering the Ingenuity Pathway Knowledge Base (IPKB) as reference set. The top significant func-tions and diseases (P-value< 0.05) were obtained by comparing functions associated with the candidate genes under selection against functions associated with all genes in the reference set, using the right-tailed Fisher exact test.

## Acknowledgments

We wish to thank Hubert Levéziel (INRA, retired), Cécile Grohs (GABI, INRAE, Jouy-en-Josas, France) and Roberta Ciampolini (Università di Pisa, Pisa, Italy) for their help during DNA extraction and management of the bovine samples from Amsterdam island collected in 1992. The genotyping of individuals from the Amsterdam island cattle breed and from the Moka cattle breed was supported by the National Institute of Agronomic Research (INRA, Animal Genetics Division, France; “SNPDOM” and “PERSAFRICA” projects). The sequencing was supported by EU (FORWARD RITA DEFI-ANIMAL project, 2015-2020) and France Génomique (CEA, Evry-Courcouronnes, France; High impact project funded “CAHWA”). MGX acknowledges financial support from France Génomique National infrastructure, funded as part of “Investissement d’Avenir” program managed by Agence Nationale pour la Recherche (contract ANR-10-INBS-09). The sequencing of individuals from the Ams-terdam island cattle was supported by the Fonds de la Recherche Scientifique – FNRS (F.R.S-FNRS) under Grant J.0134.16. Tom Druet is Research Director from the F.R.S.-FNRS.

