## Supplemental Tables and Figures for "Genomic reconstruction of the successful establishment of a feralized bovine population on the subantarctic island of Amsterdam"

| ID | Collection Year | Sex | Estimated Birth year | Genotyping | Whole Genome Sequence |
|---|---|---|---|---|---|
| TAF_5610 | 1992/03 | Female | 1984 | BovineSNP50K | n.a. |
| TAF_5614 | 1992/03 | Female | 1984 | BovineSNP50K | n.a. |
| TAF_5603 | 1992/03 | Female | 1987 | BovineSNP50K | SRR25784578 |
| TAF_5604 | 1992/03 | Female | 1989 | BovineSNP50K | SRR25784577 |
| TAF_5608 | 1992/03 | Female | 1989 | BovineSNP50K | n.a. |
| TAF_5609 | 1992/03 | Female | 1985 | BovineSNP50K | SRR25784566 |
| TAF_5612 | 1992/03 | Female | 1989 | BovineSNP50K | n.a. |
| TAF_5613 | 1992/03 | Female | 1990 | BovineSNP50K | SRR25784561 |
| TAF_5617 | 1992/03 | Female | 1986 | BovineSNP50K | SRR25784560 |
| TAF_5618 | 1992/03 | Female | 1985 | BovineSNP50K | SRR25784559 |
| TAF_5619 | 1992/03 | Female | 1985 | BovineSNP50K | SRR25784558 |
| TAF_5622 | 1992/03 | Female | 1987 | BovineSNP50K | SRR25784557 |
| TAF_MN279 | 2006/03/29 | Male | 2004 | BovineSNP50K | n.a. |
| TAF_MN280 | 2006/03/29 | Male | 2000 | BovineSNP50K | n.a. |
| TAF_MN281 | 2006/03/31 | Female | 1999 | BovineSNP50K | n.a. |
| TAF_MN282 | 2006/03/31 | Male | 2004 | BovineSNP50K | n.a. |
| TAF_MN284 | 2006/04/05 | Female | 1999 | BovineSNP50K | n.a. |
| TAF_MN285 | 2006/04/05 | Female | 2003 | BovineSNP50K | n.a. |

Table S1: Description of the individual sample from the bovine population of the Amsterdam island. Birth year were approximated from individual dental age estimation. All genotyped data (including the new ones) are available from the WIDDE repository (Sempéré *et al.*, 2015) and newly generated whole genome sequence data are publicly available in the NCBI SRA repository (PRJNA1010533 project).

| Pop. | Type | % SNP mono. | Heterozygosity | $F_{IS}$ |
|---|---|---|---|---|
| TAF | | 43.3 | 0.216 [0.212;0.222] | -0.019 [-0.034;-0.004] |
| MOK | | 12.4 | 0.263 [0.259;0.267] | 0.022 [0.016;0.030] |
| LAG | AFT | 41.0 | 0.190 [0.187;0.193] | 0.041 [0.034;0.046] |
| NDA | AFT | 34.4 | 0.214 [0.211;0.216] | 0.009 [0.005;0.014] |
| SOM | AFT | 23.5 | 0.231 [0.228;0.234] | 0.041 [0.037;0.046] |
| MAY | AFZ | 34.0 | 0.204 [0.200;0.209] | 0.037 [0.033;0.044] |
| SHK | AFZ | 23.3 | 0.255 [0.251;0.258] | -0.005 [-0.010;-0.001] |
| ZFU | AFZ | 19.5 | 0.247 [0.243;0.251] | 0.003 [-0.001;0.006] |
| ZMA | AFZ | 32.6 | 0.204 [0.199;0.209] | 0.012 [0.006;0.017] |
| ABO | EUT | 12.9 | 0.300 [0.298;0.303] | -0.039 [-0.044;-0.032] |
| AUB | EUT | 10.9 | 0.307 [0.304;0.310] | 0.011 [0.006;0.016] |
| BPN | EUT | 11.4 | 0.315 [0.312;0.318] | -0.011 [-0.018;-0.004] |
| BSW | EUT | 18.2 | 0.274 [0.268;0.277] | -0.040 [-0.048;-0.031] |
| CHA | EUT | 9.79 | 0.318 [0.315;0.320] | 0.002 [-0.004;0.008] |
| GAS | EUT | 11.3 | 0.308 [0.305;0.311] | 0.002 [-0.002;0.008] |
| GNS | EUT | 17.7 | 0.283 [0.279;0.285] | 0.020 [0.012;0.030] |
| HFD | EUT | 9.55 | 0.323 [0.321;0.326] | 0.064 [0.057;0.074] |
| HOL | EUT | 9.98 | 0.316 [0.312;0.319] | -0.000 [-0.007;0.008] |
| JER | EUT | 20.4 | 0.263 [0.258;0.266] | -0.009 [-0.017;0.001] |
| JE2 | EUT | 17.8 | 0.281 [0.276;0.284] | 0.001 [-0.005;0.012] |
| LMS | EUT | 7.41 | 0.314 [0.311;0.317] | 0.004 [-0.000;0.008] |
| MAN | EUT | 8.83 | 0.300 [0.295;0.303] | -0.013 [-0.018;-0.010] |
| MAR | EUT | 10.1 | 0.318 [0.316;0.320] | -0.013 [-0.020;-0.007] |
| MON | EUT | 13.5 | 0.291 [0.289;0.294] | -0.036 [-0.043;-0.029] |
| NOR | EUT | 11.7 | 0.300 [0.297;0.303] | -0.032 [-0.037;-0.027] |
| PMT | EUT | 8.34 | 0.321 [0.319;0.324] | -0.009 [-0.013;-0.006] |
| RDB | EUT | 21.5 | 0.258 [0.254;0.261] | -0.029 [-0.039;-0.020] |
| SAL | EUT | 13.3 | 0.294 [0.290;0.296] | 0.018 [0.014;0.027] |
| TAR | EUT | 14.4 | 0.300 [0.297;0.302] | -0.007 [-0.013;-0.000] |
| VOS | EUT | 11.1 | 0.309 [0.306;0.311] | -0.012 [-0.019;-0.008] |
| GIR | ZEB | 39.6 | 0.166 [0.162;0.172] | 0.007 [0.001;0.015] |
| NEL | ZEB | 40.8 | 0.164 [0.159;0.169] | -0.019 [-0.026;-0.013] |

Table S2: Within population genetic diversity estimated on the W50K data set. The table report for each population i) the percentage of monomorphic SNPs; ii) the average heterozygosity estimated with `poolfstat` (Gautier *et al.*, 2022); and iii) the $F_{IS}$. For the two latter statistics 95% confidence intervals derived from the block-jackknife estimates of their standard errors are given into brackets. It is important to note that heterozygosity must be interpreted with cautious and are a poor estimator of genetic diversity particularly for ZEB, AFZ and AFT breeds because of the strong SNP ascertainment bias towards SNPs overrepresented in the EUT populations in the BovineSNP50K genotyping assay (Gautier *et al.*, 2010; Matukumalli *et al.*, 2009).

| RGID | Breed Name | Sexe | SRA Run ID | Reference | Instrument | Mean Realized Coverage | |
|---|---|---|---|---|---|---|---|
| | | | | | | Auto. | X |
| TAF_5603 | AmsterdamIsland | Female | SRR25784578 | thisstudy | HiSeqX | 15.2 | 15.9 |
| TAF_5604 | AmsterdamIsland | Female | SRR25784577 | thisstudy | HiSeqX | 15.8 | 16.2 |
| TAF_5609 | AmsterdamIsland | Female | SRR25784566 | thisstudy | HiSeqX | 14.3 | 14.7 |
| TAF_5613 | AmsterdamIsland | Female | SRR25784561 | thisstudy | HiSeqX | 14.6 | 15.2 |
| TAF_5617 | AmsterdamIsland | Female | SRR25784560 | thisstudy | HiSeqX | 19.3 | 19.8 |
| TAF_5618 | AmsterdamIsland | Female | SRR25784559 | thisstudy | HiSeqX | 14.9 | 15.5 |
| TAF_5619 | AmsterdamIsland | Female | SRR25784558 | thisstudy | HiSeqX | 17.8 | 18.4 |
| TAF_5622 | AmsterdamIsland | Female | SRR25784557 | thisstudy | HiSeqX | 17.0 | 17.9 |
| ZMA_1570 | MadagascarZebu | Female | SRR25784556 | thisstudy | HiSeq2500 | 6.04 | 6.21 |
| ZMA_1583 | MadagascarZebu | Female | SRR25784555 | thisstudy | NovaSeq | 23.8 | 23.4 |
| ZMA_1585 | MadagascarZebu | Female | SRR25784576 | thisstudy | HiSeq2500 | 10.0 | 10.4 |
| ZMA_1595 | MadagascarZebu | Female | SRR25784575 | thisstudy | HiSeq2500 | 8.00 | 8.42 |
| ZMA_1649 | MadagascarZebu | Female | SRR25784574 | thisstudy | HiSeq2500 | 7.92 | 7.98 |
| ZMA_1650 | MadagascarZebu | Female | SRR25784573 | thisstudy | NovaSeq | 33.3 | 33.0 |
| ZMA_1661 | MadagascarZebu | Female | SRR25784572 | thisstudy | NovaSeq | 24.5 | 23.6 |
| ZMA_1666 | MadagascarZebu | Female | SRR25784571 | thisstudy | HiSeq2500 | 9.16 | 9.54 |
| MAY_011 | MayotteZebu | Female | SRR25784570 | thisstudy | NovaSeq | 16.1 | 15.4 |
| MAY_015 | MayotteZebu | Female | SRR25784569 | thisstudy | NovaSeq | 16.0 | 15.2 |
| MAY_021 | MayotteZebu | Male | SRR25784568 | thisstudy | NovaSeq | 17.3 | 8.86 |
| MAY_071 | MayotteZebu | Male | SRR25784567 | thisstudy | NovaSeq | 14.0 | 7.55 |
| MAY_095 | MayotteZebu | Male | SRR25784565 | thisstudy | NovaSeq | 18.6 | 9.38 |
| MAY_175 | MayotteZebu | Male | SRR25784564 | thisstudy | NovaSeq | 17.7 | 8.69 |
| MAY_219 | MayotteZebu | Female | SRR25784563 | thisstudy | NovaSeq | 13.9 | 14.0 |
| MAY_371 | MayotteZebu | Female | SRR25784562 | thisstudy | NovaSeq | 17.5 | 16.6 |
| HOL_0007 | Holstein | Male | SRR1262661 | (Daetwyler *et al.*, 2014) | HiSeq2000 | 12.1 | 2.75 |
| HOL_0018 | Holstein | Male | SRR1262672 | (Daetwyler *et al.*, 2014) | HiSeq2000 | 11.7 | 6.10 |
| HOL_0025 | Holstein | Male | SRR1262679 | (Daetwyler *et al.*, 2014) | HiSeq2000 | 11.3 | 6.00 |
| HOL_0064 | Holstein | Male | SRR1262718 | (Daetwyler *et al.*, 2014) | HiSeq2000 | 11.0 | 5.93 |
| HOL_0083 | Holstein | Male | SRR1262737 | (Daetwyler *et al.*, 2014) | HiSeq2000 | 10.6 | 4.92 |
| HOL_0117 | Holstein | Male | SRR1262771 | (Daetwyler *et al.*, 2014) | HiSeq2000 | 8.58 | 4.85 |
| HOL_0123 | Holstein | Male | SRR1262777 | (Daetwyler *et al.*, 2014) | HiSeq2000 | 8.32 | 4.76 |
| HOL_0127 | Holstein | Male | SRR1262781 | (Daetwyler *et al.*, 2014) | HiSeq2000 | 11.6 | 3.73 |
| JER_0003 | Jersey | Male | SRR1262791 | (Daetwyler *et al.*, 2014) | HiSeq2000 | 8.35 | 3.91 |
| JER_0005 | Jersey | Male | SRR1262793 | (Daetwyler *et al.*, 2014) | HiSeq2000 | 11.9 | 5.76 |
| JER_0009 | Jersey | Male | SRR1262797 | (Daetwyler *et al.*, 2014) | HiSeq2000 | 9.21 | 5.36 |
| JER_0010 | Jersey | Male | SRR1262798 | (Daetwyler *et al.*, 2014) | HiSeq2000 | 10.5 | 2.78 |
| JER_0011 | Jersey | Male | SRR1262799 | (Daetwyler *et al.*, 2014) | HiSeq2000 | 10.5 | 3.98 |
| JER_0013 | Jersey | Male | SRR1262801 | (Daetwyler *et al.*, 2014) | HiSeq2000 | 11.6 | 2.33 |
| JER_0014 | Jersey | Male | SRR1262802 | (Daetwyler *et al.*, 2014) | HiSeq2000 | 10.3 | 4.68 |
| JER_0015 | Jersey | Male | SRR1262803 | (Daetwyler *et al.*, 2014) | HiSeq2000 | 10.6 | 5.79 |
| BBI_001 | Bison bison | U | SRR6448737→740 | (Wu *et al.*, 2019) | HiSeq2000 | 25.9 | 30.7 |
| BBO_011 | Bison bonasus | U | SRR6448682→684,SRR6448670 | (Wu *et al.*, 2019) | HiSeq2000 | 29.5 | 29.5 |
| BGA_001 | Bos gaurus | U | SRR6448732→735 | (Wu *et al.*, 2019) | HiSeq2000 | 17.0 | 16.6 |
| BJA_006 | Bos javanicus | U | SRR6448720 | (Wu *et al.*, 2019) | HiSeq2000 | 16.9 | 9.26 |

Table S3: Description of the whole genome sequences, including the 24 newly generated ones, used in the study. Mean coverage for autosomes and X chromosome were estimated with *Samtools* (v1.9) *stats* program (Li *et al.*, 2009) after read duplicates removal.

| $f_3$ triplet | Estimate (x$10^3$) | 95% Confidence Interval |
|---|---|---|
| TAF;ZMA,JER | 46.35 | [43.30;49.19] |
| TAF;ZMA,JE2 | 46.43 | [43.40;49.32] |
| TAF;ZMA,HFD | 46.41 | [43.68;49.25] |
| TAF;ZMA,BPN | 46.70 | [43.84;49.35] |
| TAF;ZMA,AUB | 47.26 | [44.26;49.83] |
| TAF;ZMA,MAR | 47.25 | [44.24;50.05] |
| TAF;ZMA,SAL | 47.44 | [44.35;50.17] |
| TAF;ZMA,MAN | 47.41 | [44.47;50.15] |
| TAF;ZMA,GNS | 47.31 | [44.44;50.28] |
| TAF;ZMA,CHA | 47.65 | [44.78;50.34] |
| TAF;ZMA,NOR | 47.66 | [44.67;50.53] |
| TAF;ZMA,TAR | 47.93 | [44.77;50.46] |
| TAF;ZMA,LMS | 47.85 | [44.88;50.43] |
| TAF;ZMA,GAS | 48.24 | [45.10;50.77] |
| TAF;ZMA,HOL | 48.11 | [45.01;50.86] |
| TAF;ZMA,PMT | 48.17 | [45.21;50.76] |
| TAF;ZMA,VOS | 48.54 | [45.42;51.14] |
| TAF;ZMA,BSW | 48.48 | [45.40;51.32] |
| TAF;ZMA,ABO | 48.49 | [45.62;51.30] |
| TAF;ZMA,MON | 48.70 | [45.59;51.41] |
| TAF;ZMA,RDB | 48.71 | [45.68;51.77] |

Table S4: Estimated $F_3$ values in increasing order for all (TAF;ZMA,EUT) configurations where. The EUT population the closest to the original European source population of the TAF is expected to display the lowest value (see Figure S6.

| $f_4$ quadruplet | Estimate (x$10^3$) | 95% Confidence Interval |
|---|---|---|
| JE2,NDA;TAF,GIR | 14.28 | [12.33;16.11] |
| BPN,NDA;TAF,GIR | 14.04 | [12.33;15.90] |
| AUB,NDA;TAF,GIR | 13.98 | [12.38;15.83] |
| JER,NDA;TAF,GIR | 14.15 | [12.16;16.00] |
| SAL,NDA;TAF,GIR | 13.82 | [12.30;15.59] |
| MAR,NDA;TAF,GIR | 13.88 | [12.23;15.53] |
| GNS,NDA;TAF,GIR | 13.78 | [11.98;15.48] |
| HFD,NDA;TAF,GIR | 13.76 | [11.86;15.44] |
| NOR,NDA;TAF,GIR | 13.59 | [11.94;15.33] |
| MAN,NDA;TAF,GIR | 13.60 | [11.81;15.44] |
| TAR,NDA;TAF,GIR | 13.16 | [11.73;15.09] |
| CHA,NDA;TAF,GIR | 13.36 | [11.69;15.09] |
| LMS,NDA;TAF,GIR | 13.18 | [11.59;14.98] |
| GAS,NDA;TAF,GIR | 12.85 | [11.41;14.74] |
| PMT,NDA;TAF,GIR | 12.91 | [11.42;14.61] |
| RDB,NDA;TAF,GIR | 13.01 | [11.20;14.59] |
| HOL,NDA;TAF,GIR | 12.61 | [11.02;14.45] |
| VOS,NDA;TAF,GIR | 12.53 | [11.04;14.38] |
| ABO,NDA;TAF,GIR | 12.69 | [10.92;14.50] |
| MON,NDA;TAF,GIR | 12.50 | [10.84;14.44] |
| BSW,NDA;TAF,GIR | 12.42 | [10.49;14.33] |

Table S5: $F_4$ estimated values for all (EUT,NDA;TAF,GIR) configurations where NDA and GIR are representative of AFT and ZEB ancestry. The EUT population the closest to the original European source population of the TAF is expected to display the highest value (see Figure S6).

| Class | TAF | HOL | JER | MAY | ZMA |
|---|---|---|---|---|---|
| S | 38,024 (0.42;1.00) | 34,943 (0.41;0.98) | 30,527 (0.41;0.98) | 67,176 (0.41;0.98) | 69,283 (0.41 ; 0.99) |
| NS | 23,590 (0.26;1.00) | 24,523 (0.29;1.10) | 21,483 (0.29;1.11) | 40,266 (0.25;0.95) | 41,245 (0.25;0.95) |
| Tol. NS | 17,796 (0.2;1.00) | 18,101 (0.21;1.08) | 15,796 (0.21;1.08) | 29,801 (0.18;0.93) | 30,460 (0.18;0.93) |
| Del. NS | 4,896 (0.05;1.00) | 5,636 (0.07;1.22) | 4,960 (0.07;1.23) | 8,896 (0.05;1.01) | 9,160 (0.05;1.01) |
| LoF | 524 (0.01;1.00) | 538 (0.01;1.09) | 552 (0.01;1.28) | 996 (0.01;1.06) | 1,014 (0.01;1.05) |
| ALL | 9,036,293 | 8,520,472 | 7,440,903 | 16,223,315 | 16,706,828 |

Table S6: Number of segregating sites observed in the different populations in WGS data. The table gives the total number of variant sites and for each functional classes (S=Synonymous; NS=Non-Synonymous; Tol. NS= Tolerated NS; Del NS=Deleterious NS; and LoF=Loss-of-Function). For the four functional classes, the percentage with respect to the overall number of sites and the proportions relative to that observed in TAF population are given in parentheses.

| Class | TAF | HOL | JER | MAY | ZMA |
|---|---|---|---|---|---|
| S | 8,882 (0.41;1.00) | 11,994 (0.44;1.07) | 12,807 (0.44;1.05) | 2,078 (0.41;1.00) | 1,765 (0.39;0.94) |
| NS | 4,505 (0.21;1.00) | 5,962 (0.22;1.05) | 6,402 (0.22;1.04) | 1,126 (0.22;1.07) | 1,041 (0.23;1.09) |
| Tol. NS | 3,895 (0.18;1.00) | 5,209 (0.19;1.06) | 5,598 (0.19;1.05) | 910 (0.18;1.00) | 864 (0.19;1.05) |
| Del. NS | 399 (0.02;1.00) | 454 (0.02;0.90) | 497 (0.02;0.91) | 152 (0.03;1.63) | 129 (0.03;1.53) |
| LoF | 104 (0.005;1.00) | 132 (0.005;1.00) | 135 (0.005;0.95) | 15 (0.003;0.62) | 15 (0.003;0.68) |
| ALL | 2,150,813 | 2,719,983 | 2,939,594 | 504,199 | 455,553 |

Table S7: Number of fixed sites observed in the different populations in WGS data. Sites with non-polarized variants or found monomorphic over all samples were discarded from the counts. The table gives the total number of fixed sites and for each functional classes (S=Synonymous; NS=Non-Synonymous; Tol. NS= Tolerated NS; Del NS=Deleterious NS; and LoF=Loss-of-Function). For the five functional classes, the percentage with respect to the overall number of sites and the proportions relative to that observed in TAF population are given parentheses.

| uni. Rsb Test | Win. Pos. (chr:start-end in kb) | Win Size in kb (Nsnp) | Peak Pos. in kb (Stat. Value) | Nearest Gene (Pos. in kb ; dist. from peak) |
|---|---|---|---|---|
| JER/ZMA | 03:49,466-49,768 | 302.5 (110) | 49,482 (2.86) | ABCA4 (49,387-49,531 ; 0) |
| JER/ZMA$^\star$ | 06:85,793-88,358 | 2,564.6 (568) | 86,797 (4.39) | SLC4A4 (86,449-86,813 ; 0) |
| JER/ZMA | 06:68,561-69,197 | 636.7 (221) | 68,750 (3.96) | SCFD2 (68,539-68,936 ; 0) |
| JER/ZMA | 06:88,952-89,553 | 600.6 (168) | 88,975 ( 3.5) | LOC100847175 (88,955-88,963 ; 12) |
| JER/ZMA | 07:46,931-48,975 | 2,044 (541) | 47,277 (4.48) | SLC25A48 (47,258-47,310 ; 0) |
| JER/ZMA | 07:27,021-27,474 | 453.2 (103) | 27,056 (3.81) | MARCHF3 (26,948-27,104 ; 0) |
| JER/ZMA | 18:8,977-9,993 | 1,016 (344) | 9,074 (5.32) | CDH13 (9,138-10,154 ; 64) |
| JER/ZMA$^\star$ | 20:24,413-27,203 | 2,790 (558) | 26,070 (3.68) | ITGA2 (25,972-26,081 ; 0) |
| JER/ZMA | 20:28,067-29,177 | 1,109 (223) | 28,255 (4.57) | PARP8 (28,299-28,496 ; 44) |
| JER/ZMA | 20:21,648-22,143 | 494.6 (135) | 21,758 (4.19) | ACTBL2 (21,798-21,800 ; 40) |
| JER/ZMA | 20:22,690-23,008 | 318.7 (107) | 22,767 (4.32) | LOC112443061 (22,781-22,781 ; 14) |
| ZMA/JER | 02:5,002-5,344 | 342.5 (80) | 5,006 (3.55) | IWS1 (5,012-5,062 ; 7) |
| ZMA/JER | 12:35,363-39,777 | 4,414 (731) | 37,844 (4.01) | LOC132346771 (37,569-37,575 ; 269) |
| ZMA/JER | 12:34,871-35,327 | 455.8 (77) | 35,222 (3.69) | LOC101902228 (35,266-35,314 ; 45) |
| ZMA/JER | 17:63,890-64,781 | 890.9 (208) | 63,895 ( 4.2) | ACACB (63,846-63,961 ; 0) |

Table S8: Description of the regions harboring footprints of selection based on the $Rsb_{JER/ZMA}$ tests. Tests were carried unilaterally to provide insights into the origin of the signals. The two regions that overlap with those identified in Table 2 highlighted with a $\star$.

| Category | $-\log_{10}(P)$ range | Molecules |
|---|---|---|
| Nervous System Development and Function | 4.5;1.4 | ADGRG6,AGL,BLOC1S5,GCM1,MAPK10,SYNE2,USH1C,UTRN |
| Neurological Disease | 4.5;1.3 | ADGRG6,GCM1,HSPB3,MAPK10,OR8B8,PKHD1, SLC36A3,SYNE2,SYT16,TRPS1,USH1C,UTRN |
| Organismal Injury and Abnormalities | 4.5;1.3 | ADGRG6,AGL,BLOC1S5,BRINP2,GCM1,HSPB3,MAPK10,OR8B8, PKHD1,SLC36A3,STXBP6,SYNE2,SYT16,TRPS1,USH1C,UTRN |
| Cell Morphology | 4.2;1.4 | ADGRG6,AGL,MAPK10,PKHD1,SYNE2,TRPS1,USH1C,UTRN |
| Tissue Morphology | 4.2;1.4 | ADGRG6,AGL,MAPK10,PKHD1,SYNE2,TRPS1,USH1C,UTRN |
| Cancer | 3.7;1.3 | ADGRG6,AGL,BLOC1S5,BRINP2,GCM1,HSPB3,MAPK10,OR8B8, PKHD1,SLC36A3,STXBP6,SYNE2,SYT16,TRPS1,USH1C,UTRN |
| Gastrointestinal Disease | 3.7;1.3 | AGL,BRINP2,HSPB3,MAPK10,OR8B8, PKHD1,SLC36A3,STXBP6,SYNE2,SYT16,TRPS1,USH1C,UTRN |
| Respiratory Disease | 3.7;1.3 | ADGRG6,BRINP2,GCM1,MAPK10,OR8B8, PKHD1,STXBP6,SYNE2,SYT16,TRPS1,USH1C,UTRN |
| Cell-To-Cell Signaling and Interaction | 3.4;1.3 | GCM1,MAPK10,PKHD1,STXBP6,SYNE2,USH1C,UTRN |
| Cellular Assembly and Organization | 3.4;1.3 | ADGRG6,BLOC1S5,GCM1,MAPK10,PKHD1,SYNE2,USH1C,UTRN |
| Connective Tissue Disorders | 3.2;1.3 | ADGRG6,AGL,GCM1,TRPS1,UTRN |
| Skeletal and Muscular Disorders | 3.2;1.3 | ADGRG6,AGL,GCM1,HSPB3,STXBP6,SYNE2,TRPS1,UTRN |
| Auditory Disease | 3.2;1.5 | USH1C |
| Cardiovascular System Development and Function | 3.2;1.3 | PKHD1,SYNE2,UTRN |
| Cell Death and Survival | 3.2;1.4 | MAPK10,TRPS1,UTRN |
| Dermatological Diseases and Conditions | 3.2;1.4 | ADGRG6,BRINP2,MAPK10,OR8B8,PKHD1,SYNE2,SYT16,TRPS1,USH1C |
| Developmental Disorder | 3.2;1.3 | ADGRG6,AGL,BLOC1S5,GCM1,PKHD1,SYNE2,TRPS1,UTRN |
| Digestive System Development and Function | 3.2;1.4 | AGL,PKHD1,TRPS1,USH1C |
| Embryonic Development | 3.2;1.5 | BLOC1S5,GCM1,PKHD1,TRPS1,USH1C,UTRN |
| Hepatic System Disease | 3.2;1.6 | AGL,BRINP2,MAPK10,PKHD1,SYNE2 |
| Hereditary Disorder | 3.2;1.4 | ADGRG6,AGL,BLOC1S5,HSPB3,MAPK10,PKHD1,SYNE2,TRPS1,USH1C,UTRN |
| Metabolic Disease | 3.2;1.3 | AGL,BLOC1S5,MAPK10,PKHD1,SLC36A3,SYT16,TRPS1,UTRN |
| Ophthalmic Disease | 3.2;2 | BLOC1S5,USH1C |
| Organ Development | 3.2;1.5 | BLOC1S5,PKHD1,TRPS1,USH1C,UTRN |
| Organismal Development | 3.2;1.3 | ADGRG6,AGL,BLOC1S5,GCM1,PKHD1,SYNE2,TRPS1,USH1C,UTRN |
| Skeletal and Muscular System Development and Function | 3.2;1.5 | ADGRG6,TRPS1,UTRN |
| Tissue Development | 3.2;1.5 | ADGRG6,BLOC1S5,GCM1,PKHD1,TRPS1,USH1C,UTRN |
| Auditory and Vestibular System Development and Function | 2.9;1.7 | BLOC1S5,USH1C |
| Behavior | 2.9;1.5 | TRPS1,USH1C |
| Carbohydrate Metabolism | 2.9;1.6 | AGL |
| Organ Morphology | 2.9;1.3 | AGL,BLOC1S5,PKHD1,SYNE2,TRPS1,USH1C,UTRN |
| Renal and Urological Disease | 2.9;1.3 | AGL,PKHD1,SYNE2,SYT16,TRPS1,USH1C,UTRN |
| Cellular Function and Maintenance | 2.7;1.4 | ADGRG6,BLOC1S5,PKHD1,STXBP6,SYNE2,USH1C,UTRN |
| Cellular Movement | 2.7;1.6 | BLOC1S5,MAPK10,SYNE2,UTRN |
| Endocrine System Disorders | 2.7;1.3 | AGL,BRINP2,MAPK10,PKHD1,SLC36A3,SYNE2,SYT16,TRPS1,UTRN |
| Lipid Metabolism | 2.7;2.7 | UTRN |
| Reproductive System Disease | 2.7;1.3 | ADGRG6,AGL,BLOC1S5,BRINP2,HSPB3,MAPK10,OR8B8,PKHD1, STXBP6,SYNE2,SYT16,TRPS1,USH1C,UTRN |
| Small Molecule Biochemistry | 2.7;1.7 | ADGRG6,MAPK10,PKHD1,STXBP6,UTRN |
| Hepatic System Development and Function | 2.6;2.5 | AGL,PKHD1 |
| Hematological Disease | 2.6;1.4 | AGL,BRINP2,PKHD1,SLC36A3,STXBP6,SYNE2,SYT16,TRPS1 |
| Immunological Disease | 2.6;1.6 | AGL,BRINP2,PKHD1,SLC36A3,STXBP6,SYNE2,SYT16,TRPS1 |
| Cell Signaling | 2.6;2 | ADGRG6,MAPK10,TRPS1 |
| Cellular Development | 2.6;1.5 | ADGRG6,GCM1,PKHD1,TRPS1,USH1C,UTRN |
| Cellular Growth and Proliferation | 2.6;1.5 | ADGRG6,PKHD1,TRPS1,UTRN |
| Connective Tissue Development and Function | 2.6;1.5 | PKHD1,TRPS1 |
| Post-Translational Modification | 2.6;1.6 | HSPB3,MAPK10,UTRN |
| Cellular Compromise | 2.5;2.0 | MAPK10 |
| Reproductive System Development and Function | 2.5;2.5 | GCM1 |
| Protein Synthesis | 2.3;1.5 | AGL,OFCC1 |
| Cell-mediated Immune Response | 2.3;2.3 | MAPK10 |
| Hematological System Development and Function | 2.3;1.7 | MAPK10,PKHD1,UTRN |
| Immune Cell Trafficking | 2.3;1.9 | MAPK10,UTRN |
| Inflammatory Response | 2.3;1.7 | MAPK10,UTRN |
| Inflammatory Disease | 2.2;2.2 | UTRN |
| Drug Metabolism | 2.1;1.9 | MAPK10,STXBP6 |
| Molecular Transport | 2.1;1.6 | AGL,MAPK10,STXBP6 |
| Nucleic Acid Metabolism | 2.1;2.1 | ADGRG6 |
| Renal and Urological System Development and Function | 2.1;2.0 | PKHD1 |
| Respiratory System Development and Function | 2.0;1.3 | TRPS1 |
| Hair and Skin Development and Function | 1.9;1.6 | SYNE2,TRPS1 |
| Amino Acid Metabolism | 1.7;1.7 | MAPK10 |
| Organismal Survival | 1.7;1.5 | ADGRG6,AGL,GCM1,MAPK10,SYNE2,TRPS1,UTRN |
| Cardiovascular Disease | 1.7;1.3 | SYNE2,UTRN |
| Psychological Disorders | 1.7;1.7 | MAPK10 |
| Tumor Morphology | 1.6;1.6 | MAPK10 |
| Visual System Development and Function | 1.5;1.5 | USH1C |
| Cell Cycle | 1.4;1.4 | PKHD1 |
| DNA Replication, Recombination, and Repair | 1.4;1.4 | PKHD1 |

Table S9: List of significant diseases and functions associated with the 17 candidate genes detected under selection in the cattle population from Amsterdam island. Results were generated with Ingenuity Pathway Analysis (Ingenuity Systems, Inc., 2023)

| Ingenuity Canonical Pathways | $-\log_{10}(P)$ | Molecules |
|---|---|---|
| EGR2 and SOX10-mediated initiation of Schwann cell myelination | 3.73 | ADGRG6,UTRN |
| Agrin Interactions at Neuromuscular Junction | 2.97 | MAPK10,UTRN |
| IL-6 Signaling | 2.43 | HSPB3,MAPK10 |
| SNARE Signaling Pathway | 2.39 | STXBP6,SYT16 |
| Glycogen Degradation II | 2.11 | AGL |
| Glycogen Degradation III | 2.04 | AGL |
| Docosahexaenoic Acid (DHA) Signaling | 1.88 | STXBP6,SYT16 |
| Oxytocin Signaling Pathway | 1.78 | HSPB3,MAPK10 |
| Glycogen metabolism | 1.78 | AGL |
| IL-22 Signaling | 1.78 | MAPK10 |
| IL-17A Signaling in Gastric Cells | 1.74 | MAPK10 |
| Apelin Liver Signaling Pathway | 1.73 | MAPK10 |
| Synaptogenesis Signaling Pathway | 1.69 | STXBP6,SYT16 |
| MAPK targets/ Nuclear events mediated by MAP kinases | 1.64 | MAPK10 |
| 4-1BB Signaling in T Lymphocytes | 1.63 | MAPK10 |
| Inhibition of Angiogenesis by TSP1 | 1.63 | MAPK10 |
| MAP kinase activation | 1.57 | MAPK10 |
| April Mediated Signaling | 1.54 | MAPK10 |
| B Cell Activating Factor Signaling | 1.53 | MAPK10 |
| MIF Regulation of Innate Immunity | 1.52 | MAPK10 |
| Apelin Pancreas Signaling Pathway | 1.49 | MAPK10 |
| UVC-Induced MAPK Signaling | 1.45 | MAPK10 |
| UVB-Induced MAPK Signaling | 1.43 | MAPK10 |
| Meiotic synapsis | 1.42 | SYNE2 |
| Sensory processing of sound by outer hair cells of the cochlea | 1.42 | USH1C |
| CD27 Signaling in Lymphocytes | 1.41 | MAPK10 |
| Role of IL-17A in Arthritis | 1.39 | MAPK10 |
| GADD45 Signaling | 1.38 | MAPK10 |
| PCP (Planar Cell Polarity) Pathway | 1.38 | MAPK10 |
| Activation of IRF by Cytosolic Pattern Recognition Receptors | 1.35 | MAPK10 |
| Induction of Apoptosis by HIV1 | 1.35 | MAPK10 |
| Sensory processing of sound by inner hair cells of the cochlea | 1.32 | USH1C |
| CD40 Signaling | 1.32 | MAPK10 |
| IL-17A Signaling in Airway Cells | 1.32 | MAPK10 |

Table S10: List of significant canonical pathways associated with the 17 candidate genes detected under selection in the cattle population from Amsterdam Island. Results were generated with Ingenuity Pathway Analysis (Ingenuity Systems, Inc., 2023)

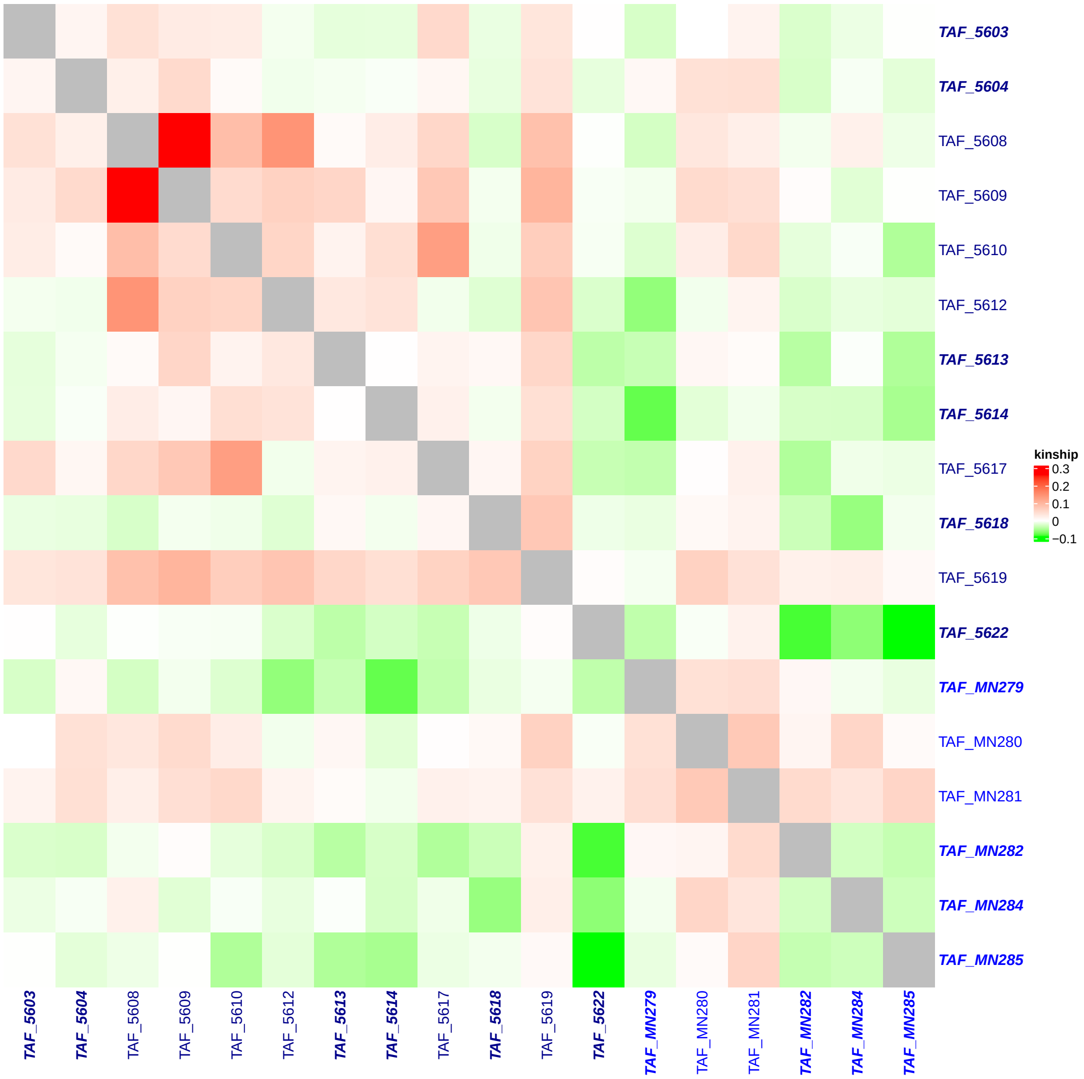


Figure S1: Heatmap of the kinship matrix among the 18 TAF individuals estimated with `King` (Manichaikul *et al.*, 2010). The names of the 12 (resp. 6) individuals collected in 1992 (resp. 2006) are colored in dark blue (resp. blue). The ten individuals deemed unrelated (up to the second degree) are in bold italic. Among the 66 pairs of 1992 samples, the estimated kinship coefficients ranged from -0.039 to 0.26 (median of 0.014). The one between TAF_5609 (born in 1985) and TAF_5608 (born in 1989) was within $[0.177, 0.354]$ suggesting a first degree relationship inferred to be a parent/offspring relationship from IBD-segment sharing (here mother/daughter from the birthdates and sex). The four coefficients between TAF_5608 and TAF_5612 ($\phi = 0.144$); TAF_5610 and TAF_5617 ($\phi = 0.131$); TAF_5609 and TAF_5619 ($\phi = 0.101$) and TAF_5608 and TAF_5610 ($\phi = 0.0887$) lie within $[0.0884, 0.177]$ suggesting a second degree relationship. Analysis of IBD-segment sharing allowed upgrading the relationship between TAF_5608 and TAF_5612 to a full-sib relationship, birthdates suggesting half-sibs or aunt-niece relationships for the three other ones. The 61 remaining coefficients between the 1992 individuals were all $< 0.0884$ (relationship more distant that second-degree). Likewise, among the 15 pairs of 2006 samples, the estimated kinship coefficients were $< 0.0884$, ranging from -0.035 to 0.074 (median of 0.013); as the 72 pairwise comparisons between individuals collected in 1992 and 2006 which ranged from -0.091 to 0.062 (median of -0.0081).

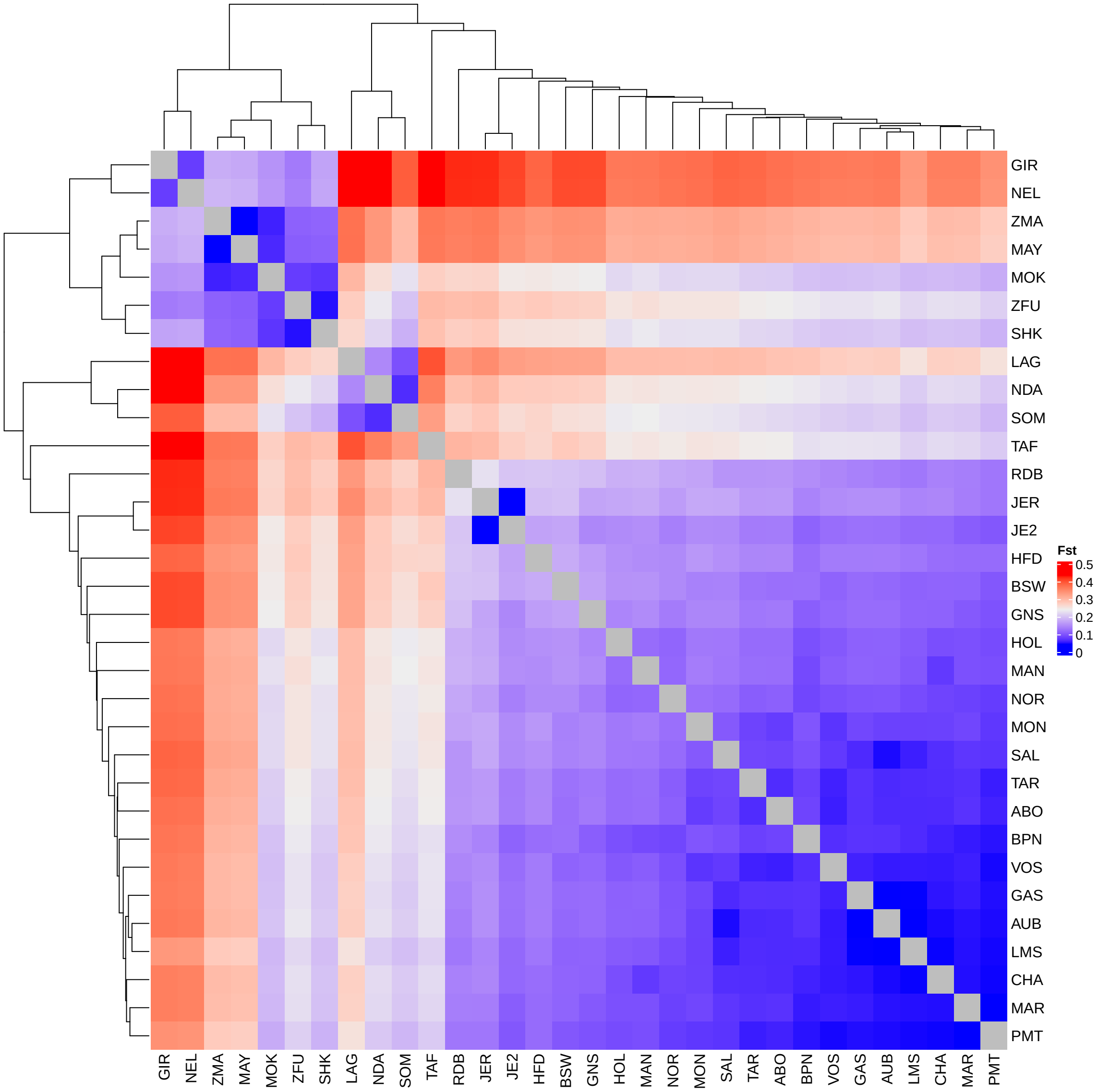


Figure S2: Heatmap of the pairwise-population $F_{ST}$ values between all the 32 populations estimated from genotyping data on the 40,426 autosomal SNPs. The estimated pairwise $F_{ST}$ values range from 0.0247 for the MAY/ZMA pair, up to 0.472 for the LAG/NEL pair, with a median of 0.203. For pairs including newly genotyped sample MOK, $F_{ST}$ values range from 0.0578 (MOK/ZMA) to 0.303 (MOK/LAG). For pairs including TAF, values range from 0.208 (TAF/PMT) to 0.436 (TAF/NEL). The TAF/PMT pairs have the lowest pairwise-$F_{ST}$ compared to others, likely because the PMT (EUT) breed has a small amount of ZEB ancestry, similar to the TAF (see Figure S4 and (Gautier *et al.*, 2010).

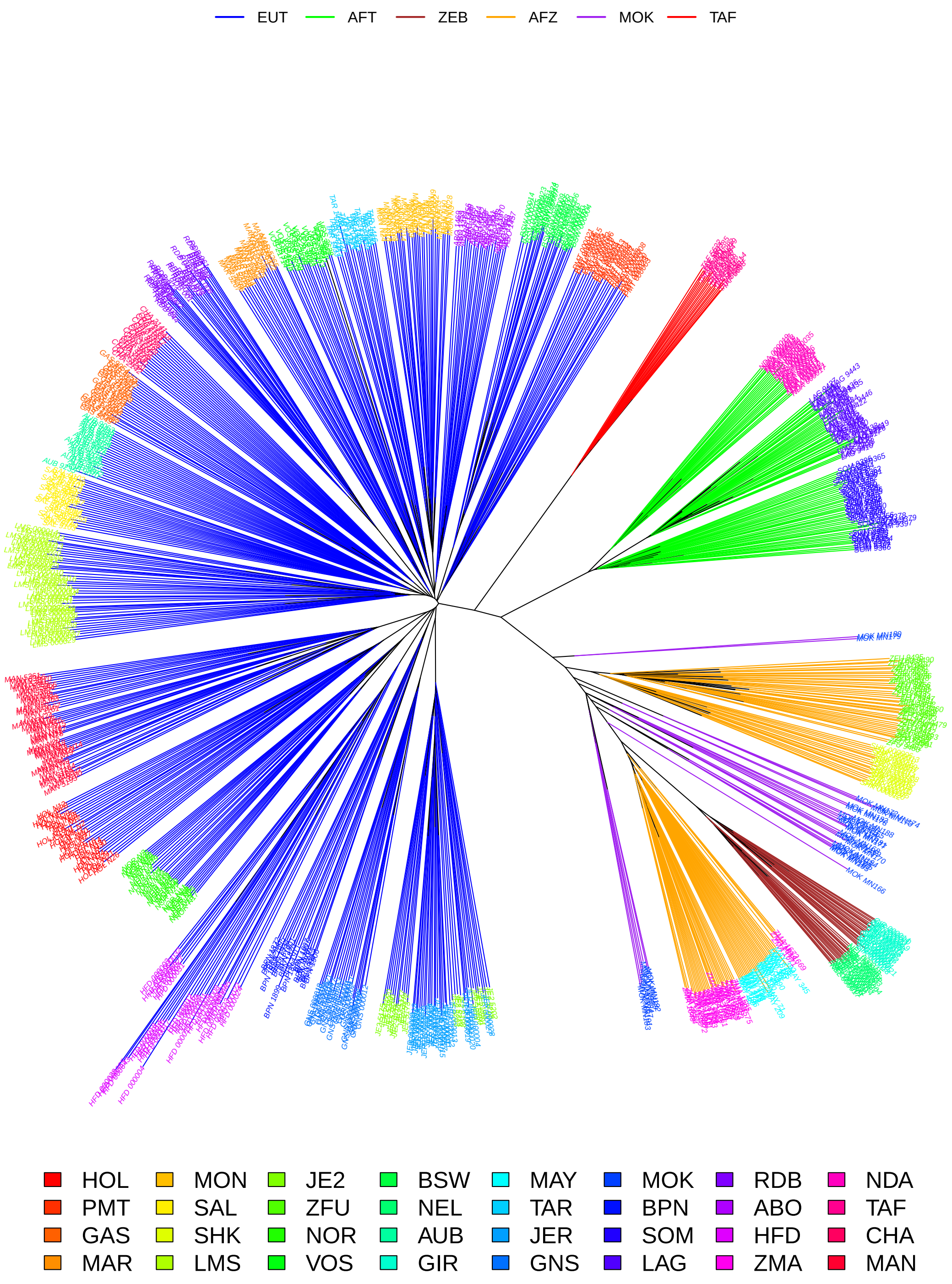


Figure S3: Neighbor-Joining tree relating the 876 individuals from the 32 populations based on allele-sharing distance computed on the 40,426 autosomal SNPs. Edges are colored according to the population group (top legend) and tip individual names according to the population of origin (bottom legend).

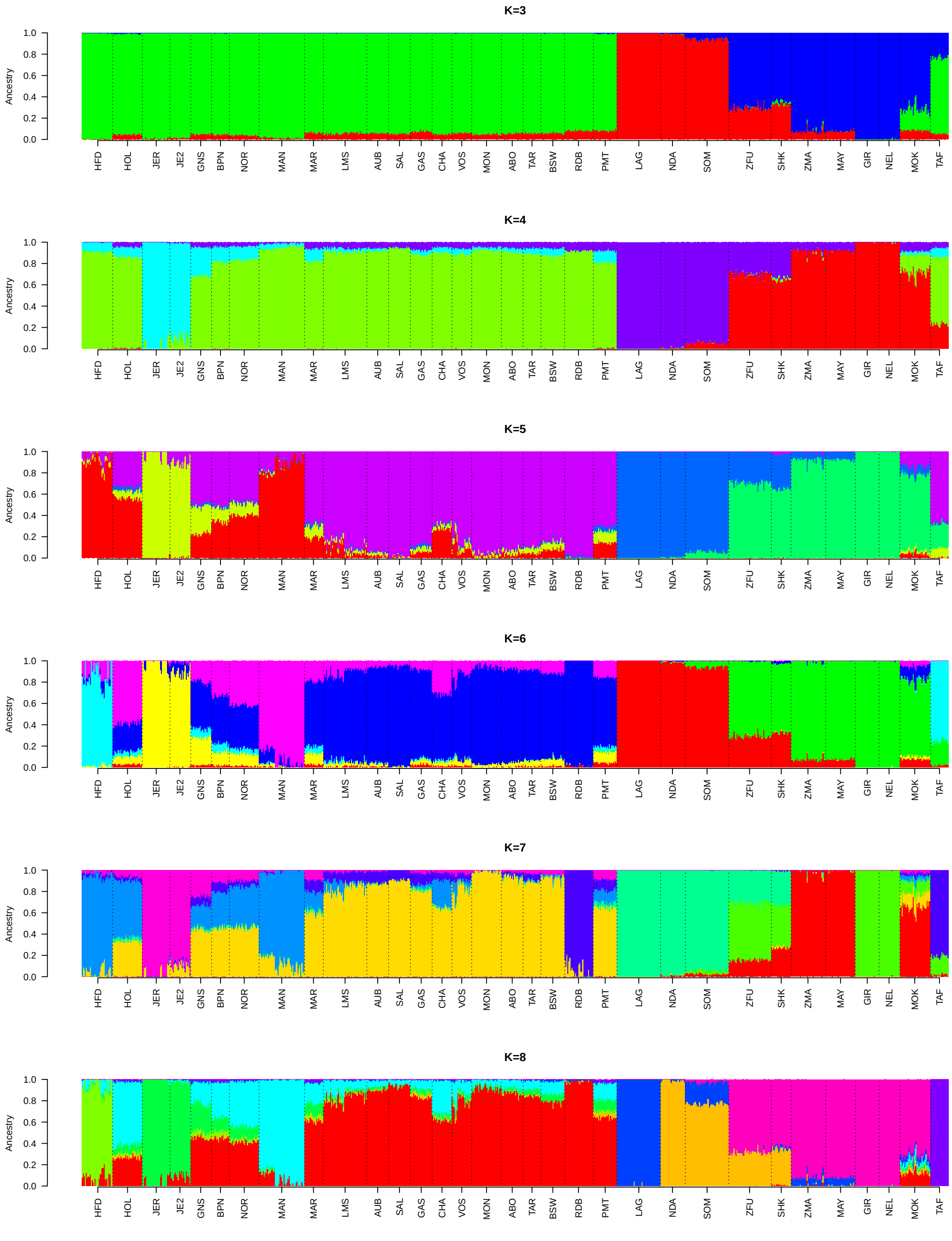


Figure S4: Unsupervised hierarchical clustering of the 876 individuals from the 32 populations based on genotyping data at 40,426 autosomal SNPs into 3 to 8 clusters.

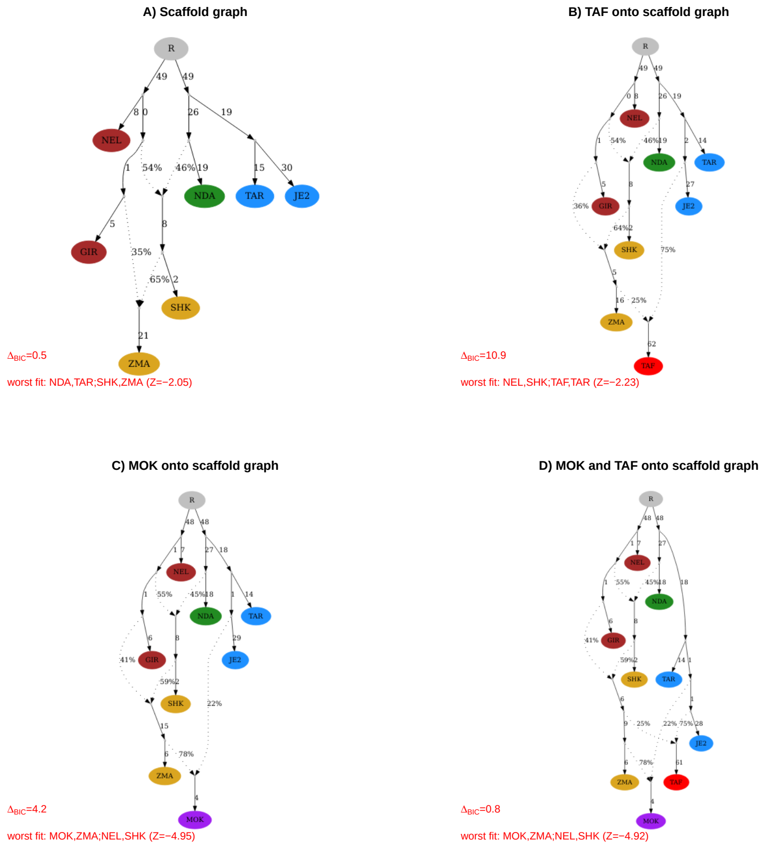


Figure S5: Admixture graph construction based on the W50K dataset with the R package `poolfstat` (Gautier *et al.*, 2022). A tree consisting of two ZEB (GIR and NEL), one AFT (NDA) and two distantly related EUT breeds (JE2 and TAR) was first used as a scaffold to which the SHK (East-African AFZ) and ZMA were jointly added to represent the history of Indian Ocean Zebus (Magnier *et al.*, 2022). The resulting best fitting graph is plotted in A) and was in agreement with results obtained by (Magnier *et al.*, 2022) using BovineHD data. Yet, the positioning of the ancestral source population of SHK that is related to the ZEB ancestor of GIR and NEL (i.e., either on the branch leading to GIR as in A) or on the branch leading to NEL or ancestral to ZEB) lead to graph with closely related BIC. B) Best fitting admixture graph among all possible ways of positioning TAF onto the scaffold graph obtained in A. This graph showing that the TAF population has an admixed origin with two sources related to JE2 ($\alpha = 75\%$) and ZMA ($1 - \alpha = 25\%$) had a very high support ($BIC$ 10.9 units lower than the graph with the second lowest $BIC$) and good fit to the observed $f$-statistics (the worst fitted f-stats associated $|Z| = 2.23$). C) Best fitting admixture graph among all possible ways of positioning MOK onto the scaffold graph obtained in A. This graph showing that the MOK population has an admixed origin with two sources, JE2 ($\alpha = 22\%$) and ZMA ($1 - \alpha = 78\%$), had good support ($BIC$ 4.2 units lower than the graph with the second lowest $BIC$) but less optimal fit to the observed $f$-statistics (the worst fitted f-stats associated $|Z| = 4.95$). Note that the worst fitted $f$-statistics ($f_4(MOK, ZMA; NEL, SHK)$) suggests that this graph fails to capture some direct ZEB (represented by NEL) ancestry to the MOK. Such ancestry would indeed contribute positively to $f_4(MOK, ZMA; NEL, SHK)$ making the observed $f$-statistics higher than the fitted $f$-statistics (and the associated Z-score negative). Such a three-way admixture origin (EUTxZMAxZEB) of the MOK would be consistent with historical record. D) Best fitting admixture graph among all possible ways of jointly positioning TAF and MOK onto the scaffold graph obtained in A. This graph is in agreement with B) and C) and suggests that MOK might not be viewed as the closest proxy to the source populations from which TAF originates although MOK and TAF share closely related ancestral EUT sources (the relative positioning of which led to graphs with close BIC).

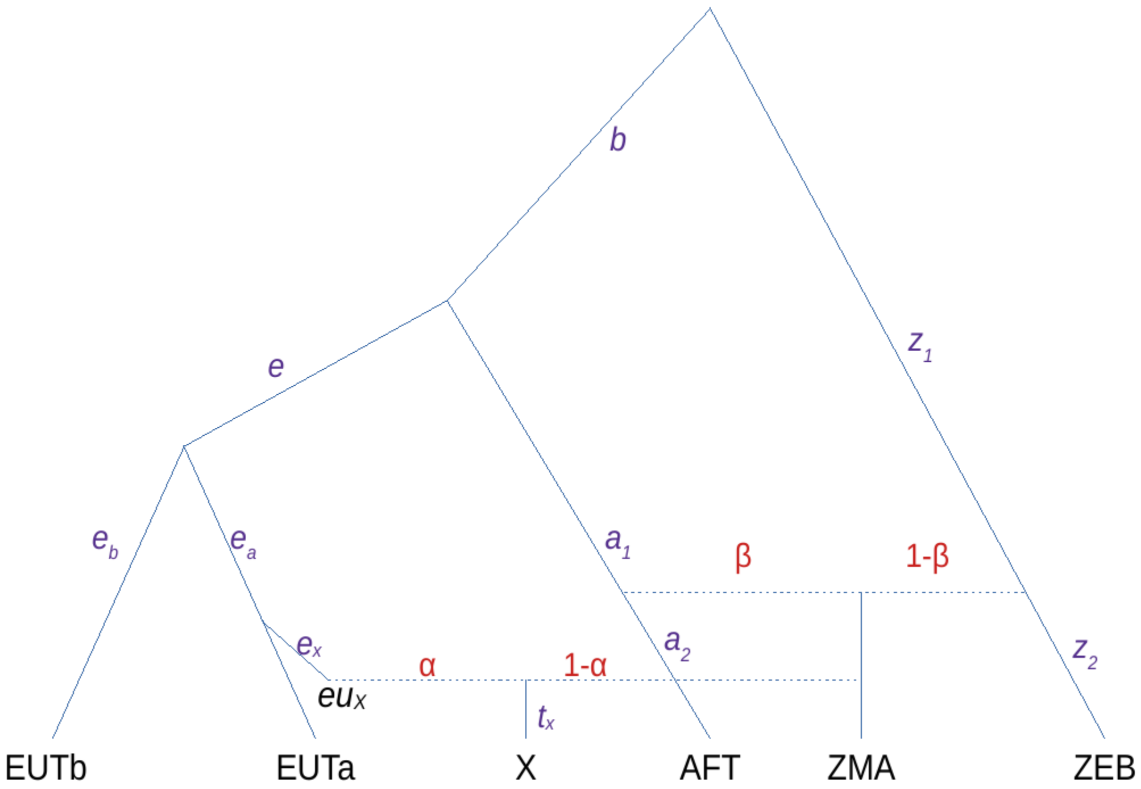


Figure S6: Schematic representation of the expected $F_4$ for various configuration under the (simplified) scenario associated to the inferred admixture graph describing the origin of an admixed population X (e.g., X=TAF or X=MOK) with one source related to one EUT and the other to the ZMA which is itself admixed. Note the ZEBxAFT admixed origin of one of the two ZMA sources can be disregarded for our configurations of interest. Under this scenario:

$$F_3(X; ZMA, EUT_b) = \tau_X - \beta(1-\beta)\left((1-\alpha)^2(z_1+b) + \alpha^2 a_1 + a_2 + e\right) = \gamma$$

$$F_3(X; ZMA, EUT_a) = \gamma - \beta(1-\beta)e_a$$

Hence, the EUT population $EUT_a$, the closest to the European source of X ($eu_X$) is expected to display the lowest $f_3$ value for the $(X; ZMA, EUT)$ configurations. Likewise,

$$F_4(EUT_b, AFT; X, ZEB) = (\alpha e - (1-\alpha)\beta a_1) = \gamma$$

$$F_4(EUT_a, AFT; X, ZEB) = \alpha e_a + \gamma$$

Hence, the EUT population $EUT_a$, the closest to the European source of X ($eu_X$) is expected to display the highest $f_4$ value for the $(EUT, AFT; X, ZEB)$ configurations. In addition,

$$F_4(EUT_b, ZEB; ZMA, EUTa) = (z_1+b)(\beta-1) - e = \tau$$

$$F_4(EUT_b, ZEB; ZMA, X) = \alpha\tau$$

The $F_4$-ratio= $\frac{F_4(EUT_b,ZEB;ZMA,X)}{F_4(EUT_b,ZEB;ZMA,EUTa)}$ then provides an estimator of the European source admixture rate $\alpha$.

A)

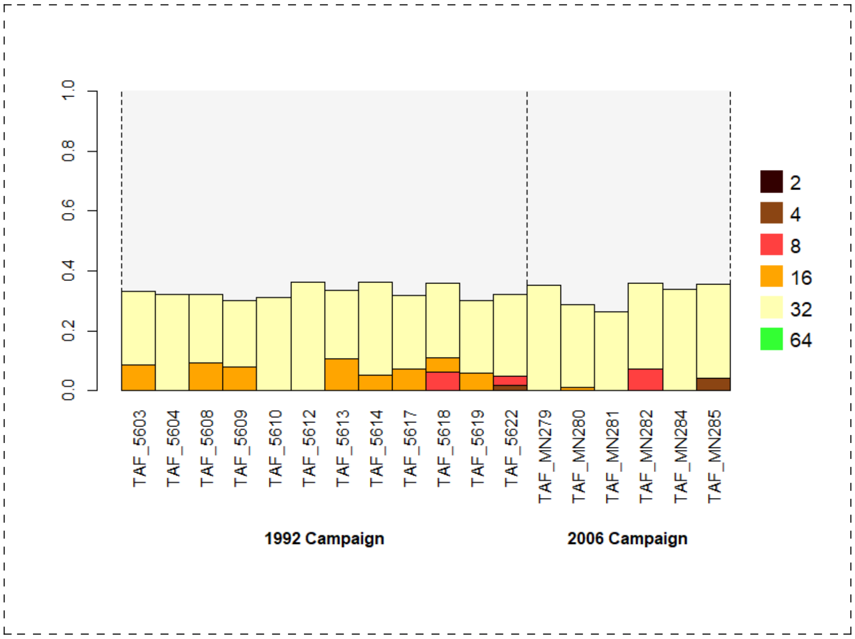


B)

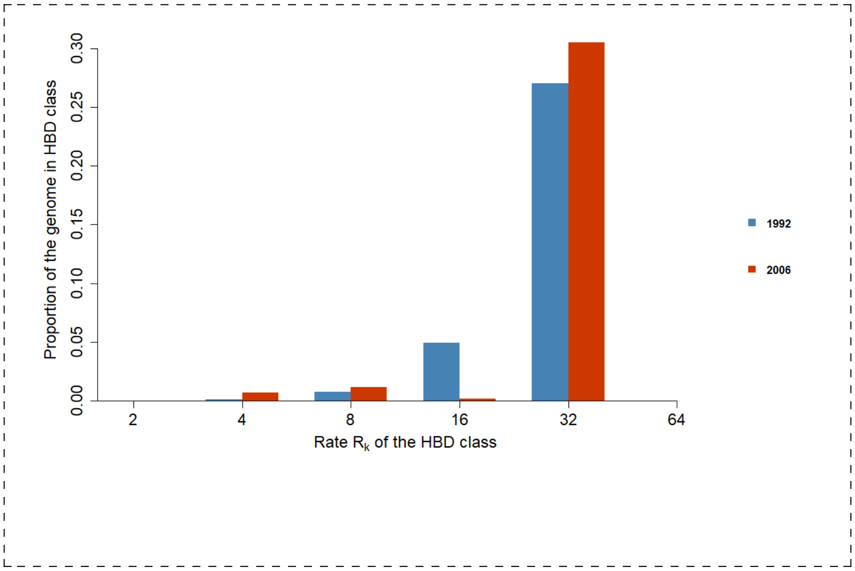


Figure S7: Partitioning of inbreeding levels in all TAF individuals using 50K SNP genotyping data (W50K dataset). A) Partitioning of the inbreeding levels into 6 different HBD classes for each individual separated by sampling campaign. B) Overall distribution of inbreeding levels among all individuals for the two sampling campaign (1992 and 2006).

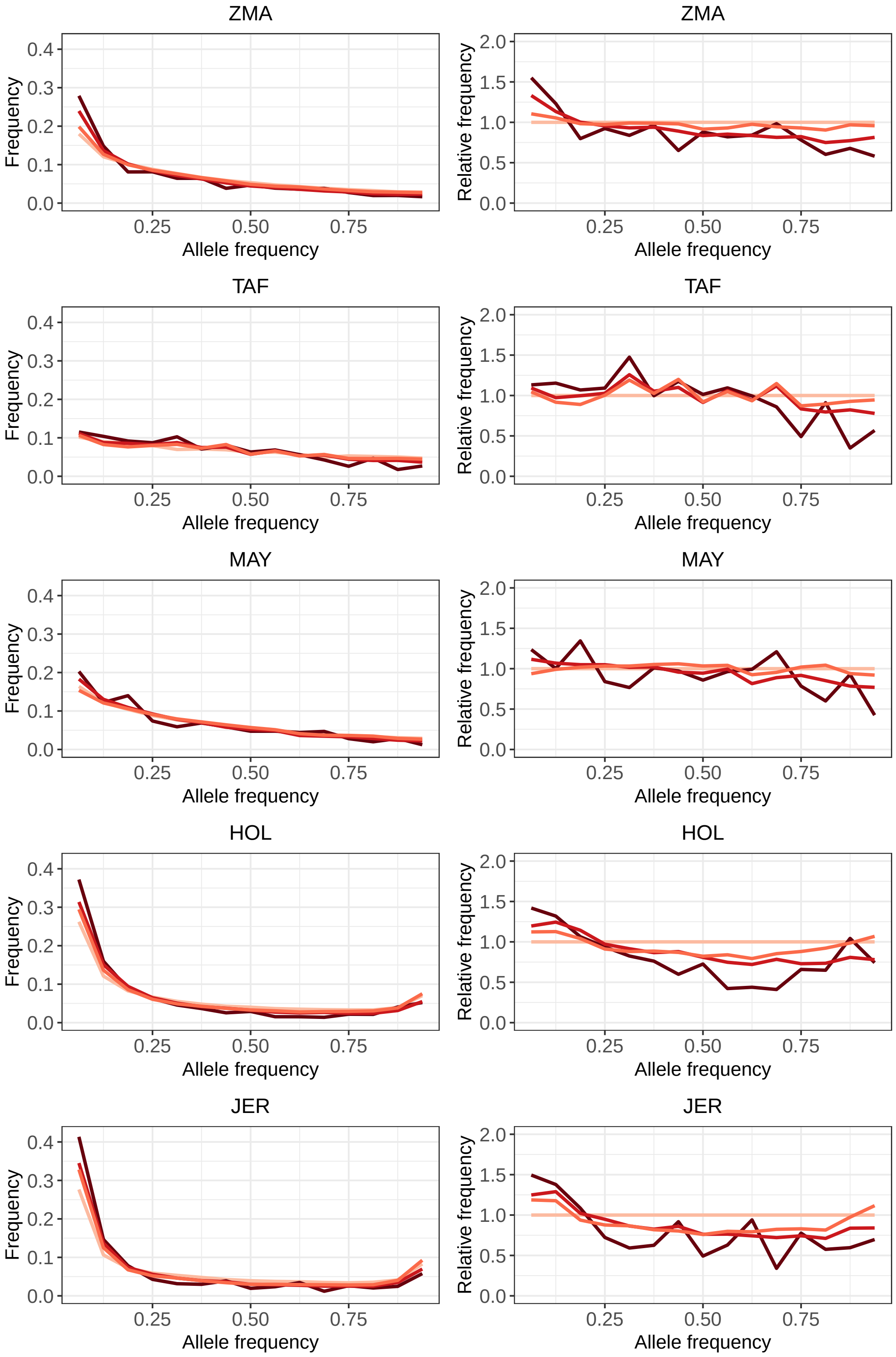




Figure S8: Comparison of the Site Frequency Spectrum (SFS) for intergenic (IG), Synonymous (S), Non-Synonymous (NS) and Loss-of-Function (LoF) variants in the five breeds colored from light to darkred as in Figure 4. The relative values were estimated by dividing with the frequencies estimated for intergenic variants.

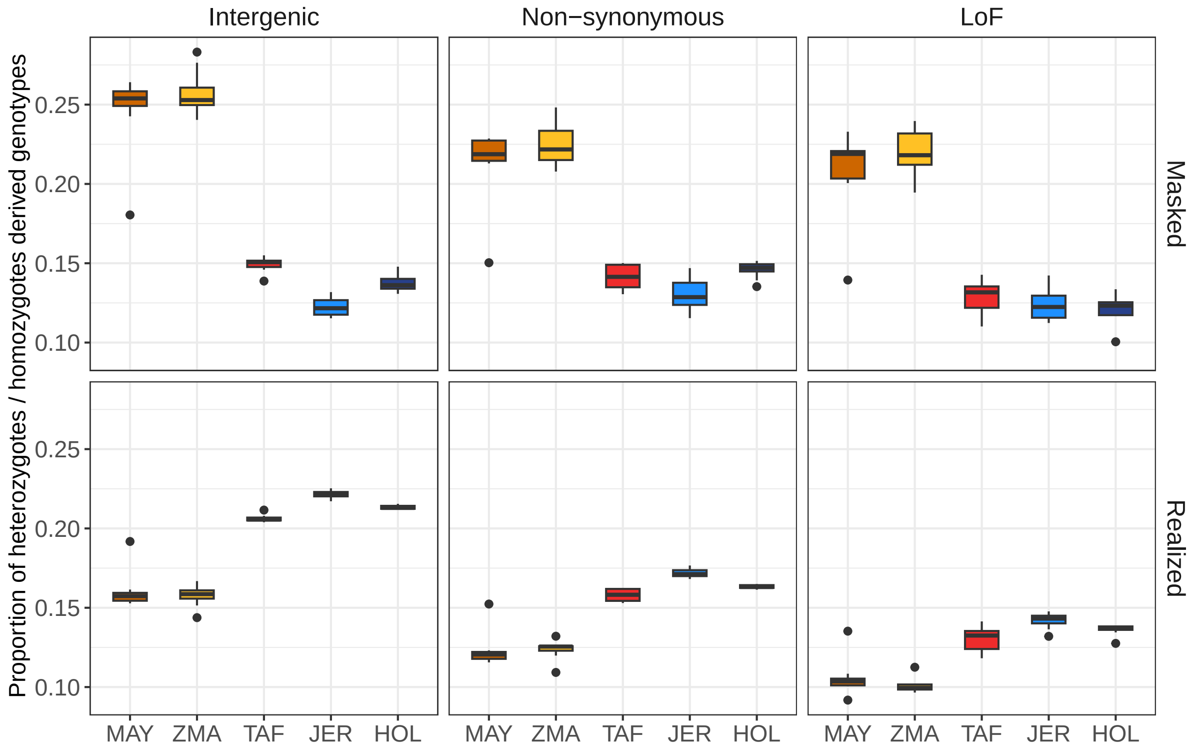


Figure S9: Distribution of the individual masked and realized loads within each population. The load were estimated per individual for intergenic, non-synonymous and loss-of-function (LoF) variants.

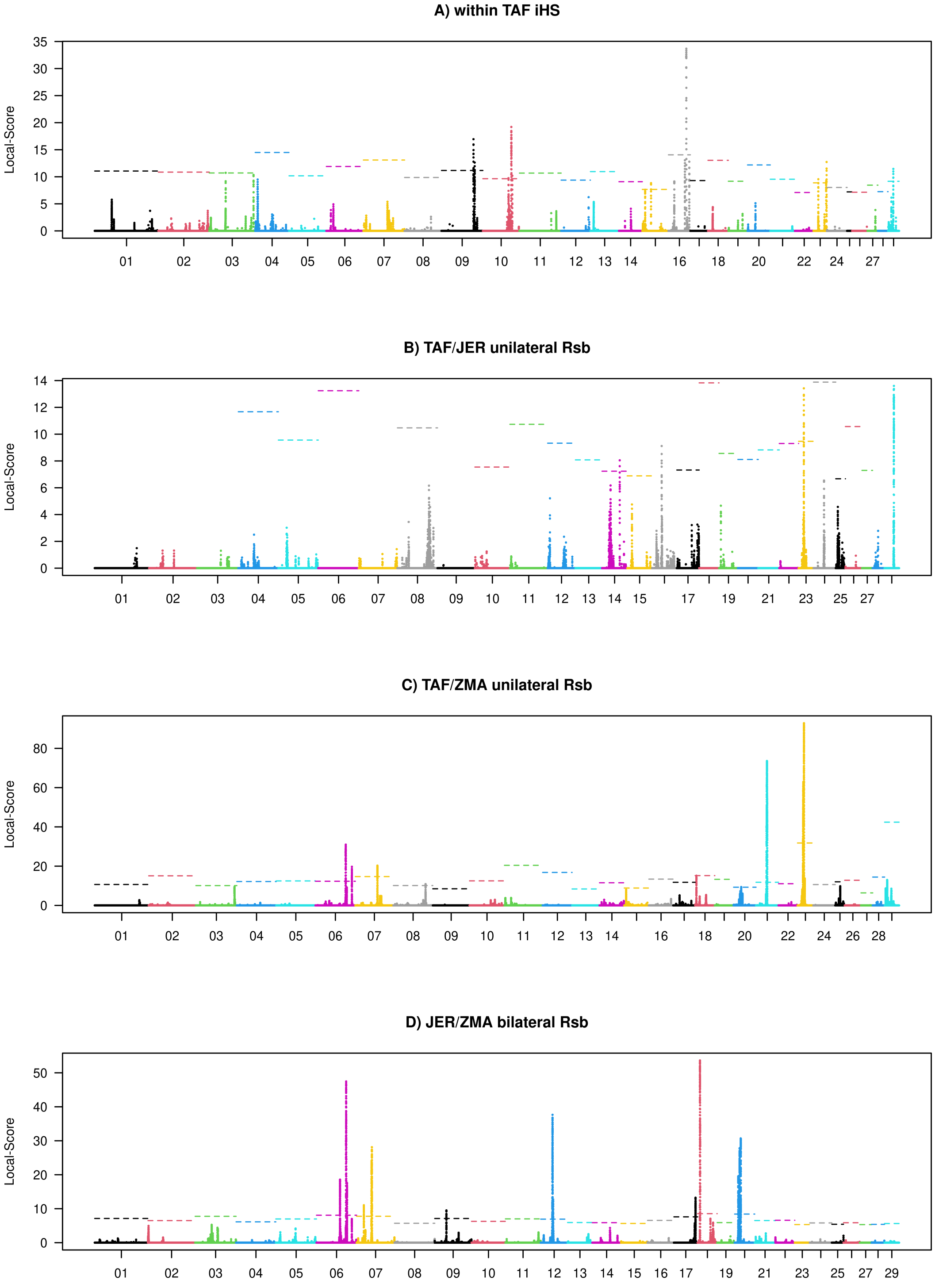


Figure S10: Manhattan plot over the bovine genome of the local scores derived from the within TAF *iHS* and the three pairwise *Rsb* statistics used to identify footprints of selection. The horizontal dashed lines indicate the chromosome specific 1% P-value threshold on the local score that was apply to select the significant windows.
